# Carbon-nitrogen metabolism drives the predatory dynamics between myxobacteria and prey bacteria by trophic interaction

**DOI:** 10.64898/2026.11.14.766615

**Authors:** Lei Zhang, Kai Yang, Guirong Yang, Lujun Zhang, Songchen Jiang, Minxin Hou, Fangyu Pang, Xianfeng Ye, Yan Huang, Zhongli Cui, Zhoukun Li

## Abstract

Interkingdom interactions represented by microbial predator and prey are crucial for soil food-web community structure and ecosystem function. However, bacteria predatory dynamics derived by metabolic interplay remain understudied. Here we examine the trophic interactions between predator myxobacteria and prey bacteria where we find that glucose input leads to predator-prey oscillations between *Corallococcus* sp. EGB and *Escherichia coli* depending on the presence of other soil microbes. Co-cultural assays revealed that glucose consumption by *E. coli* inhibits EGB predation by rapid environmental acidification via the phosphoenolpyruvate-dependent carbohydrate phosphotransferase system (PTS) and disruption of membrane homeostasis through citrate-mediated ionic chelation. In response, EGB adjusts its amino acid metabolism by producing volatile ammonia, leading to alkalinization that counteracts glucose-mediated acidification. Additionally, ammonia serves as a nitrogen source for prey bacteria growth, thereby providing a stable food source for myxobacteria by a possible farming strategy. Moreover, EGB also ingeniously exploits soil microbes *Meyerozyma guilliermondii* and *Xanthomonas* to metabolize glucose and citrate, thus adapting the predator to metabolites derived from prey in natural environments. This complex metabolite-based interplay depending on cell density and carbon-nitrogen sources leads to predator-prey oscillations between predator EGB and prey *E. coli* until a stable population equilibrium is reached. Our findings highlight the critical role of carbon-nitrogen metabolism in driving bacterial predator-prey dynamics that have implications for trophic interactions between myxobacteria and prey bacteria in soil microbial food webs.

## Introduction

The biodiversity of ecosystems forms the foundation of life on Earth and governs both ecological functions and global productivity (1, 2). As a major ecological force, predation drives species coexistence, shapes pattern of biodiversity and mediates nutrient cycling, thereby contributing to the food web complexity and population dynamics by trophic cascade effects (3, 4). Microbial food webs constitute the microscopic core of belowground ecological networks, encompassing all trophic interactions among bacteria, fungi, protists, nematodes and predatory bacteria within soil matrices (5, 6). Distinct from classical macroscopic food webs dominated by plants and animals, soil microbial food webs integrate two parallel energy pathways: the green channel fuelled by live primary producers and the brown detritus channel sustained by decaying organic matter (5, 7). Material and energy flow across multi-trophic levels via consumption, lysis and decomposition, jointly regulating soil carbon-nitrogen metabolism, nutrient mineralization and the stability of whole soil ecosystems (8). In soil microbial food webs, bacteria play a pivotal role in major global biogeochemical cycles, nutrient recycling and trophic transfer of carbon and nitrogen (9). Whereas, protists preying on bacteria and fungi could influence microbial ecosystem processes by top-down regulation (4), thereby representing a high-level of trophic regulation in the soil microbiome. Predatory bacteria such as generalist and specialized predators also possess the ability to prey on diverse soil microbes, similar to protists but with distinct predating strategies and prey ranges (10). Frequently, prey richness within microbial food web determines the performance of predator species and affects the strength of top-down control (11). However, it remains unclear how the predatory bacteria influence the composition and functional stability of prey communities by trophic interactions, and how this feeds back to population maintenance.

In bacteria, fitness, metabolic activity and population density are tightly coupled with feeding that affects predator-prey population dynamics. In microbial food webs, overexploitation from predators could lead to prey extinction (12), thereby disrupting the ecosystem. Hence, predator and prey need to keep population equilibrium by adaptive co-evolution (13), metabolic cross-talking (14), and resources availability (15). In bacteria-fungi interactions, the soil fungus *Morchella crassipes* gains fitness through the harvesting of carbon source by bacterial farming based on metabolic requirements (16). In bacteria-bacteria interactions, cell lysis through antagonism could enhance carbon flux from living bacterial cells to lysis products (17). These findings imply that microbes have evolved strategies for actively allocating resources in ecosystem, inspiring us that analogous behavior might also exist in the bacterial predation within the microbial food web, but evidence is currently lacking.

As generalist predatory bacteria found across a broad range of habitats, myxobacteria possess the capacity to feed on a vast array of bacteria and eukaryotic microbes (fungi and oomycetes) (18–22), and continue to show promise as biocontrol agents for controlling plant diseases (23–25). Predatory myxobacteria could represent a distinct module in soil microbial food webs, independent of protists and nematodes, thereby building their own trophic interactions among bacteria (6). Based on utilization of cellulose and inorganic nitrogen, myxobacteria are classified into cellulolytic or bacteriolytic groups (26). Using bacteriolytic members as models, previous studies found that myxobacteria utilize amino acids and lipids released from lysed prey as nutrients, but fail to utilize exogenous sugars, including released glucose from decomposition of *Phytophthora* cell wall (21, 27, 28). In contrast, sugars serve as an optimal carbon source for most prey bacteria (29). Since acquisition of sugar utilization would enable myxobacteria to expand its nutrition resource and environmental adaptation from an evolutionary point of view (30), it thus raises the crucial question why predatory myxobacteria fail to utilize sugars, and what is the ecological outcomes of the metabolic differences between myxobacteria and prey bacteria.

In this study, we investigate the metabolic interplay between the prominent soil inhabitants predatory myxobacteria and prey bacteria. We report that the intrinsic carbohydrate metabolism from prey and amino acid metabolism from myxobacteria drive the predatory dynamics by modulating pH values in response to glucose input, where the final population equilibrium largely depends on cell density and resource availability. We also find that myxobacteria could exploit some soil microbes to metabolize glucose and adapt glucose-mediated predation resistance. Additionally, the metabolite ammonia from myxobacteria promotes prey growth, thus suggesting a bacterial feeding behavior. Our discovery of this metabolism-driven predation dynamics adds to the knowledge of complex interplays between myxobacteria and soil microbes in microbial food webs, and opens a new avenue for understanding the ecological consequences of metabolic evolution and exact trophic positioning and functional contribution of predatory myxobacteria.

## MATERIALS AND METHODS

### Strains and growth conditions

*Corallococcus* sp. strain EGB and other myxobacterial isolates were cultured on VY/4 agar plates at 30°C for 2-3 days (31). The model strain *Myxococcus xanthus* DK1622 and related transformants were cultured on CTT medium at 30°C (32). *Escherichia coli* DH5α and other bacteria were cultured on LB plates at 37°C. *Meyerozyma guilliermondii* Y1 and other yeast species were cultured on YPD plates at 28°C. All strains and plasmids (Tables S1), primers (Table S2) used in this study are listed in the supplementary materials.

### Gnotobiotic soil co-culture assay

To determine the growth of myxobacteria and *E. coil* in the presence of glucose in soil, soil was mixed with *E. coli* at a final concentration of 5×10⁹ cells/g (cells/g soil), after which EGB were added followed by uniform soil mixing at a final concentration of 1×10⁷ cells/g. Treatments were designed as follows: Group 1,100% sterile soil; Group 2,100% non-sterile soil; Group 3, 50% non-sterile soil + 50%sterile soil; Group 4, 20% non-sterile soil +80% sterile soil; Group 5,10% non-sterile soil + 90% sterile soil; Group 6, 5% non-sterile soil + 95% sterile soil. The semi-sealed culture flasks (200 g) were incubated at 30 °C, and glucose was added at day 9, 19, and 31 to achieve a final concentration of 3.6 mg/g soil. Treatments without glucose addition were used as controls. Soil was collected and DNA extracted at different time intervals during 40 days incubation, and the biomass of EGB and *E. coli* was quantified by qPCR.

To determine the effects of cell density, glucose or casein input on the growth performance of EGB and *E. coli* in the presence of glucose in soil, micro-soil ecosystem comprising10% non-sterile soil and 90% sterile soil was designed. Treatments were designed as follows: *E. coli* was adjusted to a fixed density of 5×10^9^ cells/g soil, while the cell density of EGB was varied from 1×10⁶ to 5×10⁸ cells/g soil; EGB was adjusted to a fixed density of 1×10⁷ cells/g soil, while the inoculation level of *E. coli* was ranged from 1×10⁸ to 5×10⁸ cells/g soil; With fixed cell density of EGB (1×10⁷ cells/g soil) and *E. coli* (5×10^9^ cells/g soil), varying glucose (0.18, 0.36, 0.9, 1.8, 2.7, 3.6 mg/g soil), or casein (0, 0.5, 1, 1.5, 2, 2.5, 3, 3.5, 4, 4.5, 5 mg/g soil) with glucose (1.8 mg/g soil) were added followed by uniform soil mixing. The co-culture assays were performed in semi-sealed culture flasks and incubated at 30°C. Soil was collected and DNA extracted at different time intervals, and the biomass of EGB and *E. coli* was quantified by qPCR. The time required to achieve stable population equilibrium–defined as the absence of significant biomass fluctuations in either population—was recorded for each treatment.

### Predatory assay

Co-cultures of *E. coli* or other bacteria with different myxobacteria were carried out on solid TPM plates (10 mM Tris-HCl, 1 mM KH₂PO₄, 8 mM MgSO₄, pH 7.6) supplemented with different concentrations of monosaccharides (glucose, 1-20 mM; mannose, fructose, galactose, rhamnose, xylose, arabinose, 10 mM) and disaccharides (trehalose, maltose, sucrose, lactose, Melibiose, cellobiose, and gentiobiose, 10 mM) and casitone (1-10 g/L) when necessary. For predation assays on solid surfaces, overnight LB cultures of *E. coli* were pelleted and resuspended in liquid TPM to a final concentration of 1×10⁷ to 5×10⁹ cells/mL, 35 µL of each prey cell suspension was evenly spread onto a TPM agar plate and left to dry. EGB and other myxobacterial were cultivated, harvested, washed, and resuspended in TPM to cell density (1×10^7^ to 1×10^10^ cells/mL), and 3-µL cell suspension was spotted to the center of the prey lawn and the plates incubated at 32°C for 2-7 days. After incubation, cells were scraped from the plates for serial dilution plating to calculate the number of viable prey cells (CFU/mL), and the predation efficiency (%) was calculated as [1−( co-cultured prey cell / initial prey cell)] × 100%.

For co-cultural assays in liquid media, prepared prey bacteria (20mL, 1×10^9^ cells/mL) was transferred into TPM medium containing strain EGB with a final concentration of 1×10^7^ cells/mL, followed by culture at 30 °C for 1-2 days. The biomass of *E. coli* was quantified by dilution-plate method.

### Visual detection of extracellular pH value

To visualize extracellular acidification caused by sugar metabolism in prey bacteria, a methyl green-methyl red indicator (red at pH < 4.8, blue-green at pH 4.8-6.2, and blue-green at pH > 6.2) was incorporated into TPM agar plates containing 10 mM glucose at a final concentration of 0.002% (33). For predation assay, strain EGB with final cell density of 1 × 10⁷ cells/mL was co-cultured with different prey bacteria (5 × 10⁹ cells/mL) on TPM plate with glucose when necessary. Within 720 min incubation, color changes of the plates and corresponding pH variations and citric acid production were recorded and measured simultaneously.

### Determination of glucose utilization efficiency of bacteria

*E. coli* and the deletion strains at a cell density of 1×10⁸ cells/mL were cultivated separately in CTT and TPM liquid media containing 10 mM glucose for 48 hours, and media without glucose served as controls. Samples were collected at designated time intervals, and the supernatants were prepared by centrifugation (10,000×g for 10 min at 4 °C) and filtered through a 0.22-μm membrane filter. The residual glucose concentration in the supernatants was subsequently quantified using the DNS colorimetric method (34), and the glucose utilization efficiency (%) was calculated as: [1 - (Residual glucose content / Initial glucose content)] × 100%.

### Cell viability assay

EGB cells grown to mid-log phase were harvested, washed with TPM buffer, and the cell concentration was adjusted to 1 × 10⁸ cells/mL. The cell suspension was supplied with ethylenediaminetetraacetic acid (EDTA) or varying concentrations of citrate, and 10 mM calcium ions (Ca²⁺) and/or magnesium ions (Mg²⁺) was added in designated experimental group when necessary. The suspensions were then incubated at 30 °C for 4 h. In experiments aiming to observe the induction of cell death, propidium iodide (PI, 10 µg/mL) was added to cell suspensions and incubated for 15 minutes in the dark (35). Detailed observations of cell fluorescence were realized by a Zeiss LSM 710 confocal laser scanning microscope (Carl Zeiss, Jena, Germany and transmission electron microscopy (TEM, JEM-1400). The proportion of PI-positive (dead) cells was quantified by analyzing at least 12 randomly selected fields of view with ImageJ software.

### Synergistic predation assay of EGB and soil microbes

To examine the effects of isolated soil microbes (Tables S1) on the predation of EGB toward *E. coli* with presence of glucose, four synthetic microbial consortia were constructed with combination of *E. coli*, *Meyerozyma guilliermondii* Y1 and *Xanthomonas* sp. NJAU1. Treatments were designed as follows: *E. coli* (ConS1); *E. coli* + Y1 (ConS2); *E. coli* + NJAU1 (ConS3); *E. coli* + Y1 + NJAU1 (ConS4). The final cell concentration of *E. coli*, Y1 and NJAU1 was adjusted to 5×10⁹ cells/mL for each prey cell suspensions, and to 1×10⁷ cells/mL for EGB cell suspensions. 35μL of each prey cell suspension was evenly spread onto a TPM agar plate and left to dry. 3μL EGB cell suspension was spotted to the center of the prey lawn and the plates incubated for 3 days at 30 °C. Total DNA was extracted from cells harvested by scraping the agar surface, and the biomass of *E. coli* was quantified by qPCR, and predation efficiency (%) was calculated. During incubation, extracellular pH (at 1- and 12-h) and citrate accumulation (at 12- and 24-h) were measured as described above.

To clarify the effects of strains Y1 and NJAU1 on EGB predation, the cell-density ratios of *E. coli* to each soil microbe in ConS2 and ConS3 were systematically varied. *E. coli* in each consortium was adjusted to a final density of 5×10⁹ cells/mL, and the ratios of *E. coli* to Y1 in ConS2 and to NJAU1 in ConS3 were independently adjusted to 10:1, 5:1, 2:1, 1:1, 1:2, 1:5, and 1:10, respectively. Parallel predation experiments from the prey lawn assay were cultivated as described above. During incubation, predation efficiency (%) was calculated and extracellular pH values (at 1h) and citrate accumulation (at 12h) were investigated.

### Ammonia production in plate co-culture assay

Two-compartment petri-dish separated by a polystyrene partition was used to detect the production of volatile ammonia. On the donor side, EGB (1×10⁷ cells/mL) and *E. coli* (5×10⁹ cells/mL) were co-cultured on solid TPM or CTT medium with an initial pH of 7.6, with monoculture as a control and the petri-dishes were sealed with sealing strips. The receiver side contained blank medium with 0.002% phenol red indicator and 10 mM glucose. After incubation at 30 °C for 24-72 h, the color change of the receiver side was observed by color change visualization, and pH values and ammonia concentration were determined using the Quantofix® ammonium test kit. During 24 h incubation, EGB and *E. coli* were then co-cultured on the receiver side using prey lawn assay as described above, and the biomass of *E. coli* was measured by dilution-plate method, and predation efficiency (%) was calculated.

### Ammonia production from amino acid metabolism

To identify the amino acid metabolic pathways involved in ammonia production, EGB was inoculated into a TC3 liquid medium (TPM + 3 g/L casitone) containing different amino acids (Gly, Ser, Ala, Trp, Cys, Lys, Leu, Phe, Asp, Gln, Met, Val, Glu, Pro, His, Thr, Asn, Ile and Arg) at a final concentration of 1 g/L. Following incubation at 30°C for 48 h, the ammonia production in the culture supernatant was measured. The ammonia production of EGB in TC3 without amino acid addition was set as 100%, and the relative percentage of ammonia in treatment group was calculated.

Gene deletions in *M. xanthus* DK1622 were achieved using a double-recombination approach as described previously (36). Vector construction and transformation methods are described in Supplementary Methods. Dk1622 and derivative strains CL1019-CL1024 were cultured in CTT medium at 30 °C for 24 h, after which the ammonia level in the supernatant was measured as described above. The ammonia production of DK1622 was set as 100%, and the relative ammonia production percentages of strains CL1019-CL1024 were calculated separately.

### Growth promotion experiment of ammonia

Two-compartment petri-dish separated by a polystyrene partition was used to explore the effects of volatile ammonia on growth of soil bacteria. On the donor side, EGB (1×10⁷ cells/mL) and bacteria (*E. coli* and other soil microbes, 5×10⁹ cells/mL) were co-cultured on solid TPM or CTT medium with an initial pH of 7.6. The receiver side contained mineral salt agar medium supplemented with10 mM glucose, and different indicator bacteria (*E. coli* and other soil microbes, 1×10⁴ cells/mL) were spread-plated and incubated at 30 °C, and the petri-dishes were sealed with sealing strips. After 3 d incubation, bacterial cells from the receiving-side medium were scraped off, and the biomass of each indicator bacterium was determined using the dilution plating method. The ammonia content in the receiving-side medium was also measured by calculating the consumption rate.

Using *E. coli* as the indicator, the medium in both sides of two-compartment petri-dish was replaced with sterilized soil or pre-colonized soil (15 g per side). The receiving side (37) was incubated with *E. coli* (1×10⁴ cells/g soil) with supplement of 10 mM glucose. The donor side (right) were designed as follows: EGB, *E. coli*, EGB + *E. coli*, EGB + *E. coli* + 10 mM glucose, EGB + *E. coli* + 10 mM glucose + 3 g/L casitone. EGB and *E. coli* were used at a final concentration of 1×10⁷ cells/g soil and 5×10⁹ cells/g soil, respectively. After 10 days of co-culture, soil was collected and DNA was extracted, and the biomass of EGB and *E. coli* was quantified by qPCR. The predation efficiency in the donor side (right) and the fold change of *E. coli* biomass in the receiving side (37) were calculated.

### Statistics and reproducibility

All experiments were performed with at least three independent biological replicates, and reproducible results were consistently obtained. Data are presented as the mean ± standard error (SE). Statistical significance was assessed using one-way analysis of variance (ANOVA) followed by Student’s t-test, performed with SPSS 20.0 (IBM Inc., Chicago, IL, USA). A value of *p* < 0.01 was considered highly significant, and significant differences between groups are indicated by lowercase letters. Pearson correlation coefficients were calculated to analyze relationships between variables. All micrographs and image-based observations were independently repeated at least three times with similar results,and the numbers of “n” denotes the sample size.

## Results

### Glucose input leads to population oscillations of predator *Corallococcus* sp. EGB and prey *E. coli* in soil

To date, there is still limited experimental evidence from microbe-based systems to support the existence of trophic cascades in soil decomposer food webs dominated by bacteria. To reveal possible trophic interactions between predatory myxobacteria and prey, we established a gnotobiotic soil co-culture system composing of the predator *Coralllococcus* sp. EGB that cannot utilize glucose and prey *E. coli* capable of utilizing glucose (Fig. S1). We precisely manipulated the abundance of background microbiota by adjusting the mixing ratio of non-sterile soil to sterile soil, and investigated how the biomass of background microbiota and glucose input affect this predator–prey interactions (Fig. 1a). Here, samples were collected and DNA extracted at specific time intervals to quantify changes in the abundance of *E. coli* during the co-incubation by qPCR (Fig. 1b). The abundance of *E. coli* declined continuously from day 3 to 9 from treatments without glucose input and maintained the population stability at 1×10^5^ copy number from day 15 to 40. This population change of prey *E. coli* was also observed from high proportion of non-sterile soil (100% and 50%, groups 2 and 3) with glucose input (Fig. 1b). By contrast, glucose input increased the biomass of *E. coli* from treatments with low proportion of non-sterile soil (5%-20%, groups 4-6) to varying degrees during co-culture with predator EGB and eventually maintained the population stability at day 30 or 35 (Fig. 1b). Notably, in group 5, *E. coli* biomass increased to 1.7×10⁷ copy number following glucose addition at day 9, with a secondary peak after the second glucose addition at day 19, indicative of a biphasic fluctuation pattern (Fig. 1b). However, the *E. coli* population from treatment with 100% sterile soil reduced to 1.5×10^5^ copy number at day 9 but increased to 1.4×10⁸ copy number after glucose input and maintained this population stability from day 15 to 40 (Fig. 1b). These results show that predation of *E. coli* by EGB results in reduced biomass of prey bacteria in soil, while glucose input restores the prey *E. coli* population by possible predation resistance, and this outcome may rely on soil microbes.

**Fig. 1.**
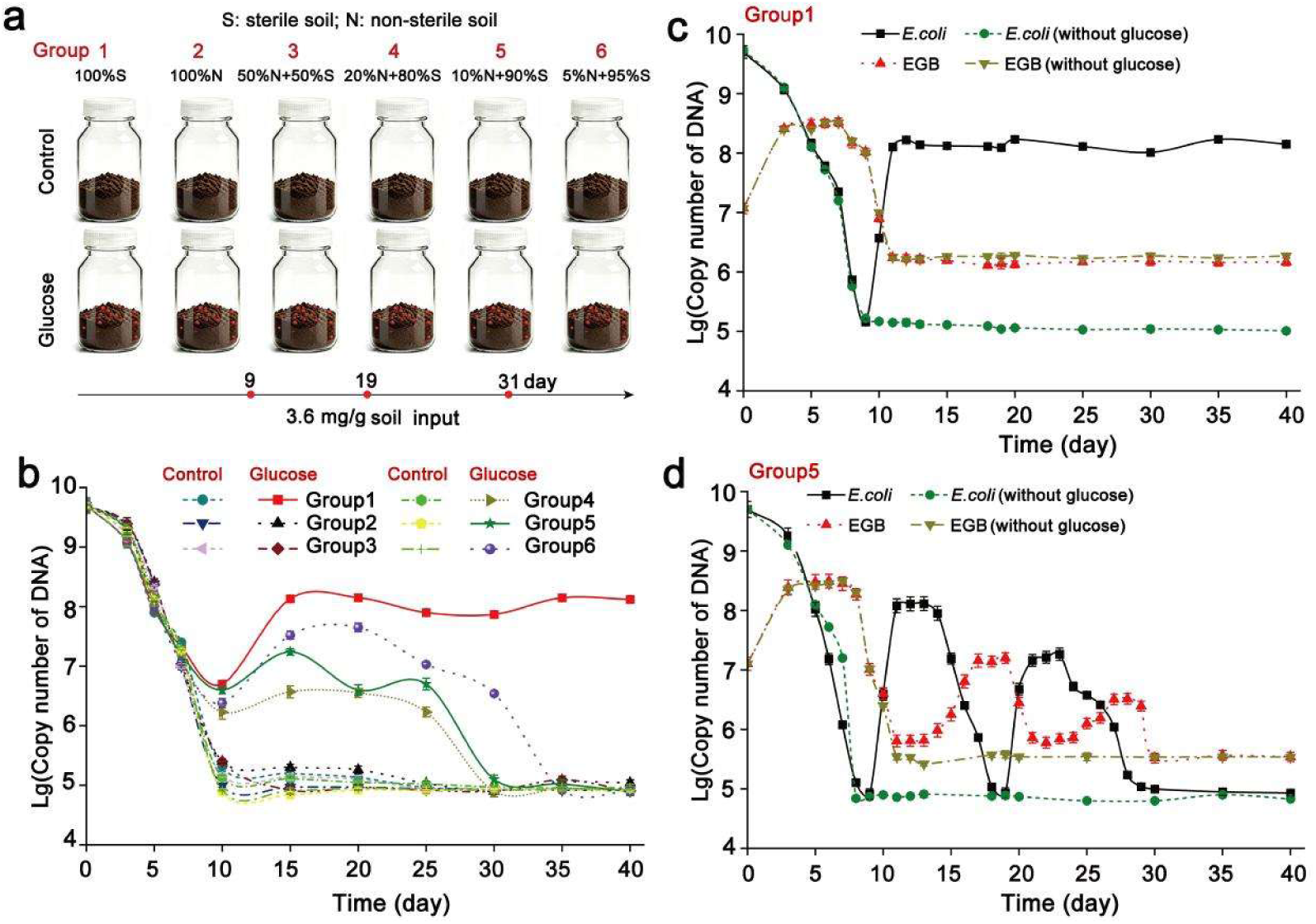
Population dynamics of *Corallococcus* sp. EGB and *E. coli* in soil in response to glucose input. **a)** Schematic of the soil co-culture system. The experiment was comprised of six treatment groups (each with twelve replicates) by changing the ratio of sterilized to non-sterilized soil. Soil co-culture experiments with mixtures of EGB (1×10⁷ cells/g soil) and *E. coli* (5×10⁹ cells/g soil) in response to glucose input (3.6 mg/g soil) at 9, 19, and 31. S denotes sterilized soil; N denotes non-sterilized soil. **b)** The cell population of *E. coli* from different treatments and times were quantified via real-time quantitative PCR (qPCR), with results expressed as copy number of DNA. Quantitative analysis of EGB and *E. coli* biomass in the 100% sterilized soil treatment **c)** and the 10% non-sterilized + 90% sterilized soil treatment **d)** from different times by qPCR. Error bars denote ± standard error of the mean (SD).

To uncover the population dynamic of predator EGB and prey *E. coli* during co-culture, we further designed the soil co-culture experiments from group 1 (100 % sterile soil) (Fig. 1c) and group 5 (10% non-sterile + 90% sterile soil) (Fig. 1d). Here, samples were collected at specific time intervals to measure changes in the abundance of EGB and *E. coli* during the co-incubation by qPCR. During the initial co-culture stage (day 0 to 9), EGB exhibits effective predation toward *E. coli*, resulting in obvious reduction in *E. coli* biomass (reduced to 1×10^5^ copy number) accompanied by a concurrent increase in EGB biomass (increased to 3×10⁸ copy number) (Figs. 1c-d). In contrast to prey *E. coli*, predator EGB biomass reduced to 4×10⁶ copy number and maintain population stability from day 10 to 50 in group 2, independent of glucose (Fig. 1b), indicating that population dynamic of predator EGB and prey *E. coli* in response to glucose input depend on soli microbes.

Whereas, in group 5 (10% non-sterile + 90% sterile soil), we found that, once *E. coli* abundance dropped below a critical threshold, EGB biomass also declined, consistent with classic predator–prey dynamics under resource limitation depending on population density. When glucose was added at day 9, the biomass of prey *E. coli* quickly increased by nearly three orders of magnitude (increased to 1 × 10^8^ copy number) and kept population stability from day 11 to day 13, suggesting that glucose support prey *E. coli* rapid growth, leading to a high stable density. However, EGB biomass remained stable without biomass change (day 11 to 13), but surged nearly 25-fold (increased to 1.6×10⁷ copy number) from day 14 to 17, coinciding with decreased *E. coli* biomass (day 14 to 18) (Fig. 1d). This population dynamics was repeatable as revealed from day 19 to 30 with two-stage glucose input at day 19, after which EGB and *E. coli* coexisted stably at a fixed ratio of 2:1 from day 31 to 40 (Fig. 1d). The experimental predator-prey cycles manifested as a regular community succession reveals the existence of predator-prey oscillations between predator EGB and prey *E. coli* derived by possible trophic interactions, thus presenting experimental demonstration of coupled oscillations in a microbial food web.

### Carbohydrates mediates the inhibitory interaction between predatory myxobacteria and prey bacteria

In the soil co-culture experiment, we found that glucose input in sterile soil triggers the explosive growth of *E. coli* while causing a stagnation of the predatory function of EGB (Fig. 1c), indicating possible predation resistance. To explored the underlying reason, the effect of glucose on myxobacteria life cycle was explored with EGB as the model (20). EGB was unable to utilize glucose (Fig. S1), and the presence of glucose exerted no effects on its growth rate, motility, EPS production, fruiting body formation or myxospore development (Figs. S2a-d). However, glucose inhibited EGB predation against *E. coli* in a dose-dependent manner, reducing predatory efficiency at 1 mM and completely abolishing it at 10 mM (Figs. 2a-b, Fig. S2e). This finding aligns with observations from sterile soil microcosms, wherein glucose input halted predatory interaction between EGB and *E. coli* (Fig. 1c).

**Fig. 2.**
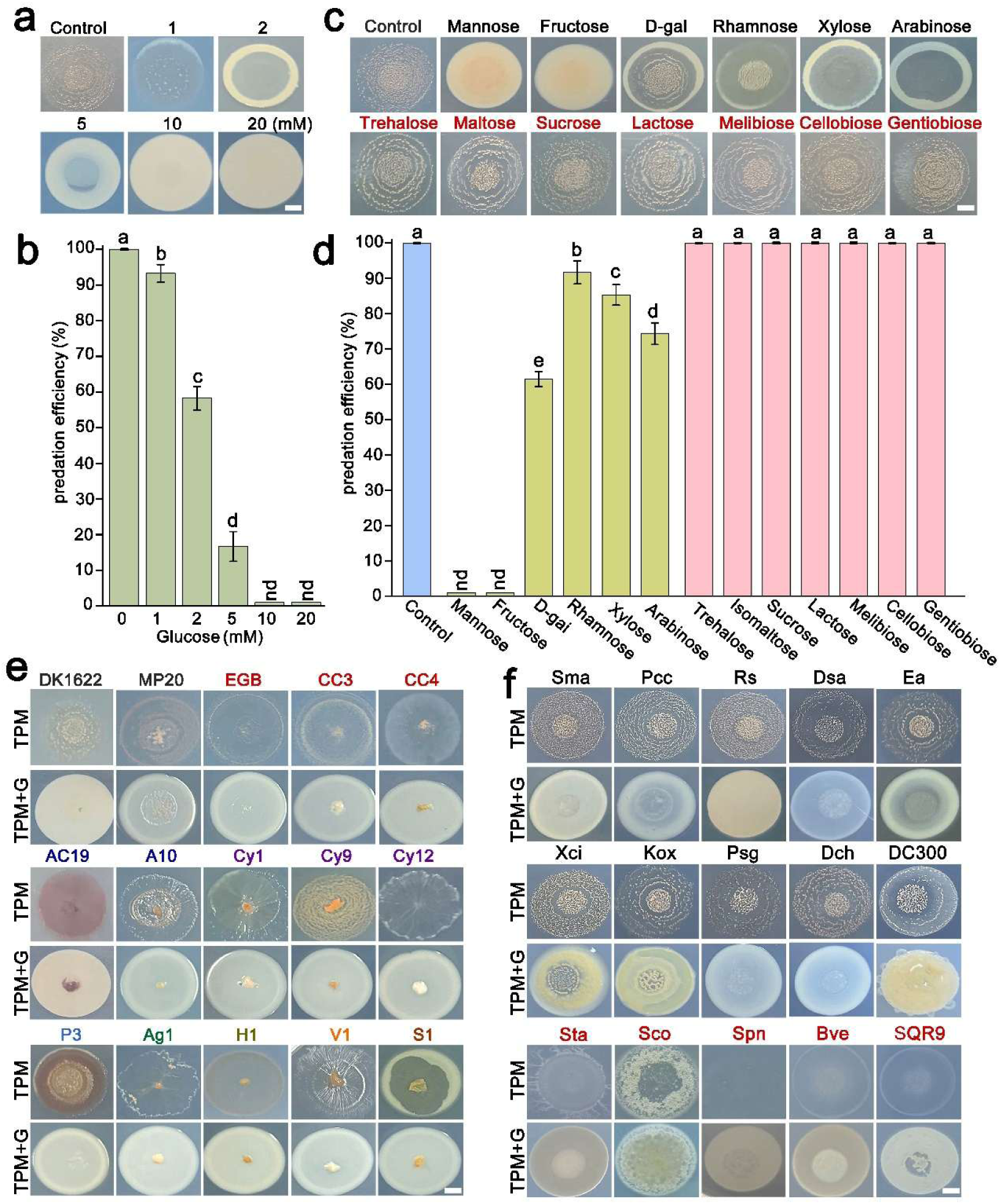
Inhibition of myxobacteria predation by bacteria with the presence of saccharides. **a, b)** Predation behavior of EGB with glucose (0-20 mM) on TPM plate (a) and the corresponding predation efficiency (b). **c, d)** Impact of different sugar types (10 mM) on predation behavior of EGB on TPM plate (c) and the corresponding predation efficiency (d). **e)** Co-culture assay of myxobacterial isolates with *E. coli* on TPM agar plates containing no glucose (top) or 10 mM glucose (bottom). DK1622: *M. xanthus*; MP20: *Myxococcus* sp.; EGB, CC3 and CC4: *Corallococcus* sp.; AC19 and A10: *Archangium* sp.; Cy1, Cy9 and Cy12: *Cystobacter* sp.; P3: *Pyxidicoccus* sp.; Ag1: *Aggregicoccus* sp.; H1: *Hyalangium minutum*; V1: *Vitiosangium cumulatum*; S1: *Stigmatella erect*. **f)** Co-culture assay of EGB with different prey bacteria on TPM agar plates containing no glucose (top) or 10 mM glucose (bottom). Strain abbreviations: *Serratia marcescens* (Sma), *Pectobacterium carotovorum* (Pcc), *Ralstonia solanacearum* (Rs), *Dickeya solani* (Dsa), *Erwinia amylovora* (Ea), *Xanthomonas citri* (Xci), *Klebsiella oxytoca* (Kox), *Pantoea agglomerans* (Psg), *Dickeya chrysanthemi* (Dch), *Pseudomonas syringae* DC300 (DC300), *Staphylococcus aureus* (Sta), *Streptomyces coelicolor* (Sco), S*treptococcus pneumoniae* (Spn), *Bacillus velezensis* (Bve), *Bacillus subtilis* SQR9 (SQR9). EGB and other myxobacterial isolates (1×10⁷ cells/mL, 3 µL) were inoculated to the center of *E. coli* and other bacteria (5×10⁹ cells/mL, 35 µL) spots on TPM agar plates with different saccharides or glucose (10 mM) and documented after 2 d. Predation efficiency was calculated by measuring the number of *E. coli* colony forming units (CFU) that determined by serial dilution and selective plating. G denotes glucose. Error bars denote ± standard error of the mean (SD), and different lowercase letters indicate statistically significant differences (*p* < 0.01, n=3) between groups based on Student’s t-test. “nd” indicates that no predation was detected. Scale bar, 2 mm.

To further elucidate the specificity of this response, we evaluated the effects of various monosaccharides and disaccharides. All tested hexose and pentose monosaccharides restrained EGB predation to varying degrees, with mannose, and fructose exerting the most pronounced inhibitory effects, akin to glucose (Figs. 2c-d). Conversely, the test disaccharides showed no effects on predation of EGB toward prey *E. coli* (Figs. 2c-d). To determine whether this predation inhibition was unique to EGB, we inoculated other predatory myxobacteria onto the center of the prey *E. coli* lawn. We found that all the 15 predatory myxobacteria isolates from 9 genera, including *Myxococcus, Corallococcus, Archangium, Cystobacter, Pyxidicoccus, Aggregicoccus, Hyalangium, Stigmatella, Vitiosaangium* failed to prey on *E. coli* with presence of 10 mM glucose (Fig. 2e), suggesting that glucose-mediated predation inhibition is wide-spread in myxobacteria taxa. We next sought to determine whether this predation inhibition could be triggered by other prey bacteria. EGB was co-cultured with diverse bacterial species on TPM agar plates, including 10 Gram-negative bacteria and 5 Gram-positive bacteria. We observed that EGB preyed on all the tested species, but not with in the presence of 10 mM glucose (Fig. 2f). These results imply that monosaccharides, exemplified by glucose, protects prey bacteria from myxobacterial predation, and this carbohydrate-mediated predation inhibition is widespread in bacterial communities.

### PTS-mediated extracellular acidification endows dominating prey resistance against predator EGB

To understand how prey bacteria inhibit myxobacteria predation with glucose, we examined the effects of carbohydrate metabolism of bacteria on predation. Given that the culture supernatant of *E. coli* from glucose-contained medium inhibited EGB predation and growth, and this inhibition could be relieved in buffered condition for Aci-SUP1 (pH 7.0) (Fig. S3), we hypothesized that acidification of the medium resulting from prey sugar metabolism contributes to the glucose-mediated inhibition of myxobacteria predation. To visualize this process, we used bromocresol green, a pH indicator that shifts from green to pink under acidic conditions. As expected, *E. coli* progressively acidified the extracellular milieu, with the indicator transitioning to blue and pink with pH values dropping to 5.1 and 3.3 from 7.6 after 1h and 12h, respectively (Fig. 3a), and the pH changes did not rely on the predator EGB (Fig. S4). Additionally, glucose-mediated extracellular acidification was also observed across diverse bacterial prey (10 Gram-negative and 5 Gram-positive bacteria) (Fig. S5), indicating that prey bacteria acidify the environment quickly by innate carbohydrate metabolism. To investigate the involvement of the phosphoenolpyruvate (PEP)-dependent carbohydrate phosphotransferase system (PTS) and glycolysis pathway that commonly participate in carbohydrate uptake and metabolism in bacteria (38), we created Δ*ptsI-ptsH* and Δ*HK* mutants of *E. coli* by deleting the genes QGF37390.1, QGF37391.1 (PTS) and QGF37413.1 (Embden–Meyerhoff–Parnas, EMP), and found that glucose utilization efficiency of the Δ*ptsI*-*ptsH* mutant was significantly impaired in nutrient-deficient conditions and markedly reduced in nutrient-rich conditions (Fig. 3b). In contrast, the Δ*HK* mutant retained glucose utilization efficiency akin to that of wild type (Fig. 3b). Consistently, the Δ*HK* mutant retained the ability to acidify the environment (Fig. 3a) and inhibit predation (Figs. 3c-d). In contrast, the Δ*ptsI-ptsH* mutant failed to acidize the extracellular environment (Fig. 3a) and significantly reduced resistance against EGB predation in solid and liquid conditions, and this inhibition could be restored by complementing individual genes (Figs. 3c-d).

**Fig. 3.**
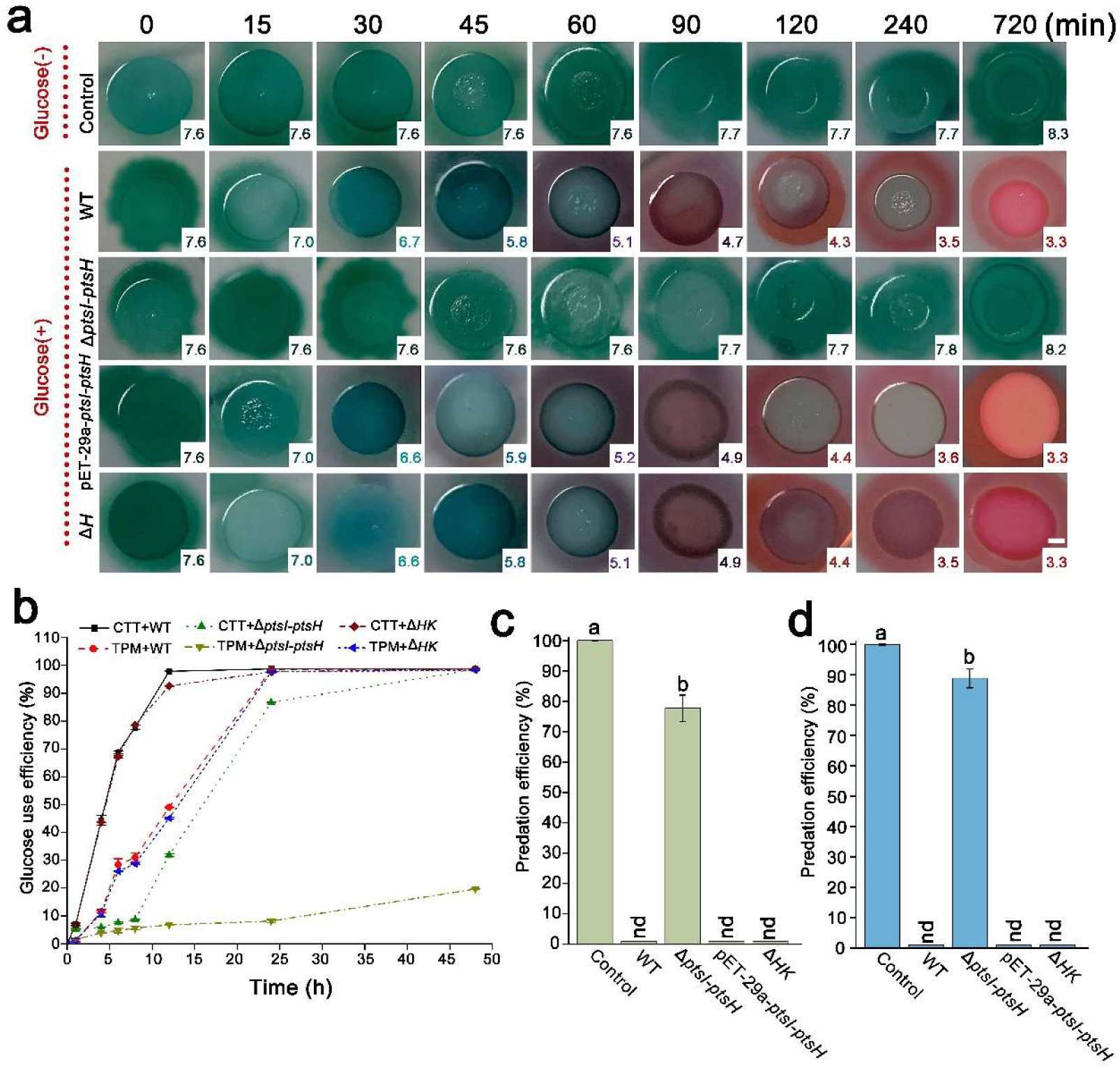
PTS-dependent extracellular acidification confers prey bacteria resistance against EGB predation. **a)** pH change illustrated by the color change of the indicator (methyl phenol green-methyl red, 0.005%). EGB (1 × 10⁷ cells/mL, 3 µL) was inoculated to the center of *E. coli* spots (5 × 10⁹ cells/mL, 35 µL) on agar plates and documented at various times until 12 h (120 min) incubation. Color changes of indicated bacterial strain lawns and their corresponding pH values were recorded at indicated time points. Scale bar, 2 mm. **b)** The effects of PTS and EMP pathway deletions on the glucose utilization capacity of *E. coli* at different growth points. Predation efficiency of EGB during co-culture with *E. coli* from TPM agar plates **(c)** and liquid medium **(d)**. *E. coli* was co-cultured with EGB with 10 mM glucose at 30°C for 2 days, quantitative analysis of the biomass of *E. coli* by colony counting (CFU), and the predation efficiency was calculated as described above. WT, wild-type *E. coli*; Δ*ptsI-ptsH*, PTS system deletion mutant; pET-29a-*ptsI*-*ptsH*, Δ*ptsI-ptsH* complementation strain; Δ*HK*, EMP pathway deletion mutant. Error bars denote ± standard error of the mean (SD), and different lowercase letters indicate statistically significant differences (*p* < 0.01, n=3) between groups based on Student’s t-test; “nd” indicates that no predation was detected. Scale bar, 2 mm.

We then wondered why myxobacteria fail to launch predation in response to glucose-mediated extracellular acidification induced by prey bacteria. We found that *E. coli* withstands a wide range of pH values (4.0-10.0) without affecting its growth (Fig. S6a). Whereas, EGB only tolerated narrow pH values (6.0-8.0), with weak growth at pH 9.0 and no growth below pH 6.0 (Fig. S6b), suggesting that EGB is more acid-sensitive than *E. coli* and thereby producing the predation inhibition. These findings indicate that prey bacteria with superior acid adaptability exploit PTS system to rapidly acidify surroundings environment in the presence of glucose, thus act a defense strategy against myxobacteria predation.

### Citrate facilitates auxiliary prey resistance by disrupting the membrane homeostasis of predator EGB

Given that Aci-SUP4 still inhibited the predation and growth of EGB even after its pH was restored to 7.0 (Fig. S3), suggested that factors other than acidification were also involved in this process. To investigate this, EGB was co-incubated with Aci-SUP1 and Aci-SUP4 in the presence of propidium iodide (PI). Aci-SUP4 induced EGB death (as indicated by PI staining) regardless of pH adjustment, but not for Aci-SUP1 (Fig. S7a). In contrast, Aci-SUP4 exhibited no cytotoxic effect on *E. coli* cells themselves (Fig. S7b). In addition, deletion of *ptsI-ptsH* reduced the prey resistance toward predator EGB but did not fully abolish it (Figs. 3c-d). Revealing that short-term exposure to an acidic environment only restricts EGB growth and predation but was insufficient to kill cells. These results suggest that non-acid factor(s) beyond acidification contribute to glucose-mediated prey defense via cell killing.

Previous studies have shown that *Serratia maurcescens* induces rapid acidification of the extracellular medium by glucose catabolism via production of lactic, acetic, pyruvic and citric acids (39). Therefore, we speculated whether these organic acids contribute to *E. coli* defense against EGB predation. As expected, all the four organic acids at a concentration of 1mM facilitate prey *E. coli* resistance against EGB predation (Fig. S8a). Unexpectedly, adjusting the pH values of citric acid (pH 4.0) to 7.0 also failed to restore the predation of EGB, in contrast to lactic, acetic, and pyruvic acids (Fig. S8b). Subsequent assays showed that citrate-mediated prey resistance was dose-dependent, with a minimum effective concentration of 0.1 mM (Fig. S8c), and co-incubation of EGB with 0.1- and 1 mM potassium citrate for 4 h resulted in 4% and 98% cell death, respectively, as indicated by PI staining and live cell counting (Fig. 4a). In contrast, citrate exhibited no cytotoxic effect on *E. coli* cells (Fig. S9a). Moreover, the lethal effect of citrate on EGB was independent of pH in the range of 6.0-8.0 (Fig. S9b) and consistent with the results in Aci-SUP4 (Fig. S7a), indicating that citrate-mediated lethal effect is not related to acidity. Furthermore, compared with the extracellular acidification mediated by PTS within 1h, citrate accumulation in Aci-SUP required more than 4 h to reach the threshold concentration of 0.1 mM (Fig. S10). These findings suggest that PTS and citrate from glucose metabolism work together to endow the antipredator defenses by two sequential actions, prey bacteria utilize the PTS system to rapidly acidify the surrounding environment and inhibits myxobacteria growth and predation, followed by the production of the intermediate metabolite citrate that kills myxobacteria at an accumulated effective concentration.

**Fig. 4.**
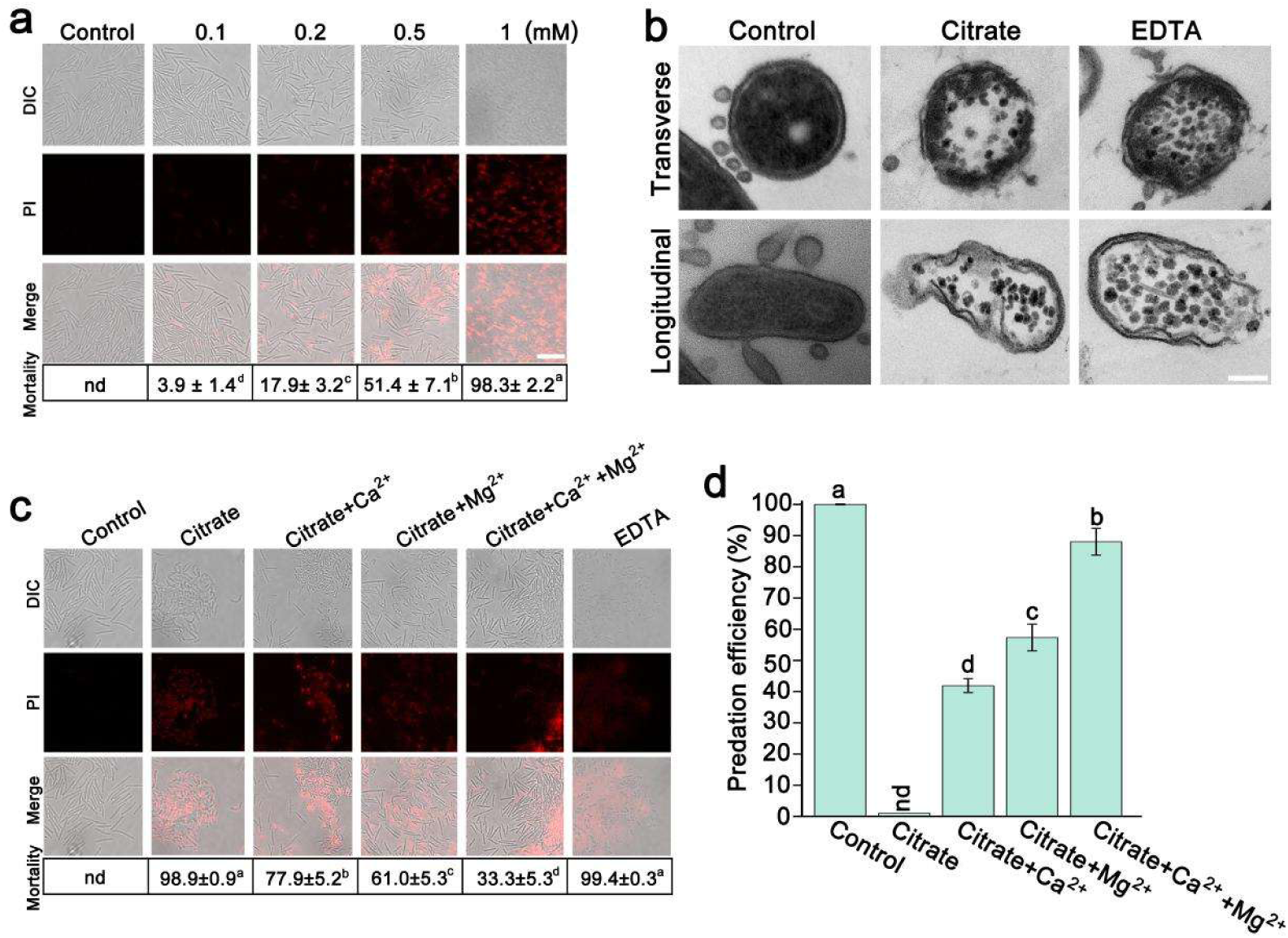
Citrate inhibits EGB by disrupting membrane integrity via cation chelation. **a)** Incubation of EGB with citrate (0-1mM) at 30°C for 4-hour. Fluorescence indicates propidium iodide (PI) staining and the induction of cell death to EGB. Cell mortality was quantified using ImageJ software. Scale bar: 5 μm. **b)** TEM analysis of the cell membrane of EGB after co-culture with citrate and EDTA (0.5 mM). Scale bar: 200 nm. **c)** Effects of Ca²⁺/Mg²⁺ supplementation (10 mM) on the cell viability of EGB after treatment with 0.5 mM citrate. Cell viability of EGB was observed by PI staining and cell mortality was quantified using ImageJ software. Scale bar: 5 μm. **d)** Effects of Ca²⁺/Mg²⁺ supplementation (10 mM) on predation efficiency of EGB during co-culture with *E. coli* on TPM plates. Calculations were performed as described above. Error bars denote ± standard error of the mean (SD), and different lowercase letters indicate statistically significant differences (*p* < 0.01, n=6) between groups based on Student’s t-test. “nd” indicates no cell death.

Studies on antimicrobial activity of sodium citrate against *Streptococcus pneumoniae* have revealed that chelating cationic ions from cell wall components by high concentrations of citrate sodium (>25 mM) can disturbed membrane function and subsequently cause cell lysis (40, 41). However, EGB exhibited a higher sensitivity to citrate. We next sought to uncover the mechanisms that underlies the citrate-mediated lethal effect. TEM observations and PI staining showed that citrate or EDTA treatment damages the cell membrane integrity of EGB, resulting in significant cell killing along with leakage of cytoplasmic components (Figs. 4b-c), whereas *E. coli* was unaffected (Figs. S9a, c). The addition of Mg^2+^ and Ca^2+^ attenuated the antibacterial effect of citrate on EGB cells, which partly recovered cell morphology and viability (Fig. 4c), and their combination reduced the lethal effect of citrate by 65.6%, thus revealing an additive effect (Fig. 4c). Cation protection was also observed in predation assays, where supplementation with Mg²⁺ and/or Ca²⁺ reversed citrate-mediated resistance (Fig. 4d). By performing a transcriptome analysis (RNA sequencing (RNA-seq)) on EGB cultured with 0.5 mM citrate, we found that citrate treatment altered the expression of 2,782 genes (Fig. S11a), including 558 that were significantly differentially expressed relative to the control. Gene Ontology (GO) analysis revealed that over half of upregulated genes (94 genes, >54%) were associated with membrane homeostasis or metal ion binding, including integral membrane components, fatty acid biosynthetic process, magnesium ion binding, calcium ion binding, and metal ion binding (Fig. S11b). These findings indicate that citrate from glucose metabolism endows prey resistance by chelating cations in the cell wall membrane of predator EGB, causing the outer membrane dysfunction and loss of viability.

### EGB exploits soil microbes-mediated carbohydrate consumption to realize predation under glucose-rich condition

Comparative predation assays in sterile versus non-sterile soils revealed opposite phenotypes upon glucose input that EGB predation was suppressed in sterile soil, but not for non-sterile soil (Figs. 1b–d), and incubation of *E. coli* in sterile soil with glucose led to rapid acidification along with substantial citrate accumulation, in contrast to that of non-sterile soil (Fig. S12), indicating a potential ecological role of soil microbes in regulating the interaction between myxobacteria and prey bacteria in response to the glucose input.

To investigate the contribution of key soil microbes, we isolated 150 bacteria and 12 yeast strains from non-sterile soil samples supplied with glucose (Table S3). Functional screening revealed that four yeast isolates— *Meyerozyma guilliermondii* Y1, *Candida tropicalis* Y2, *Pichia etchellsii* Y3, and *Pichia farinosa* Y4—exhibited efficient glucose consumption with lower levels of acidification and citrate compared to *E. coli* (Fig. S13). Among these, *M. guilliermondii* Y1 was selected as the representative strain due to its superior glucose utilization efficiency, minimal acidification and citrate production (Fig. S13). Additionally, we identified an isolate *Xanthomonas* sp. NJAU1 with resistance to EGB predation (Fig. S14) and possessed rapid and pH-independent citrate consumption (Fig. S15). Hence, we hypothesized that *M. guilliermondii* Y1 and *Xanthomonas* sp. NJAU1 alleviate glucose-induced predation inhibition through the consumption of specific carbon metabolites. We then tested bacterial predation by constructing synthetic communities comprising different combinations of EGB, *E. coli* and soli isolates with the presence of glucose (Fig. 5a). These co-culture assays were again done on TPM agar plates. We found that Y1 in ConS2 significantly promotes the predation of EGB toward *E. coli* with the presence of glucose, whereas NJAU1 and ConS3 did not (Fig. 5b). Compared to predation inhibition in ConS1 (EGB *+ E. coli*), EGB exhibited improved predation efficiency (increased to 29.8%) toward *E. coli* in ConS2 (EGB *+ E. coli*+ Y1) (Fig. 5b). Notably, although NJAU1 alone did not relieve predation inhibition, the predation efficiency increased to 97.1% in ConS4 (EGB *+ E. coli*+ Y1 + NJAU1) (Fig. 5b). The pH values in ConS1 decreased to 4.5 at 2 h and 3.4 at 24 h, while in ConS2 they were 6.7 and 6.9 and ConS4 they were 6.1 and 8.8 at the corresponding time points (Fig. 5c). Additionally, the citrate levels decreased to 0.06 mg/g in ConS2 and were undetectable in ConS4, compared to 0.22 mg/g in ConS1 after 24 h (Fig. 5d). The citrate levels of ConS3 decreased to 0.04- and 0.004-mg/g after 12 h and 24 h, respectively (Fig. 5d). These results indicate that *M. guilliermondii* Y1 could alleviate the predation inhibition by competitively utilizing glucose and consuming citrate, whereas *Xanthomonas* sp. NJAU1 facilitating myxobacteria predation by very efficiently scavenging citrate.

**Fig. 5.**
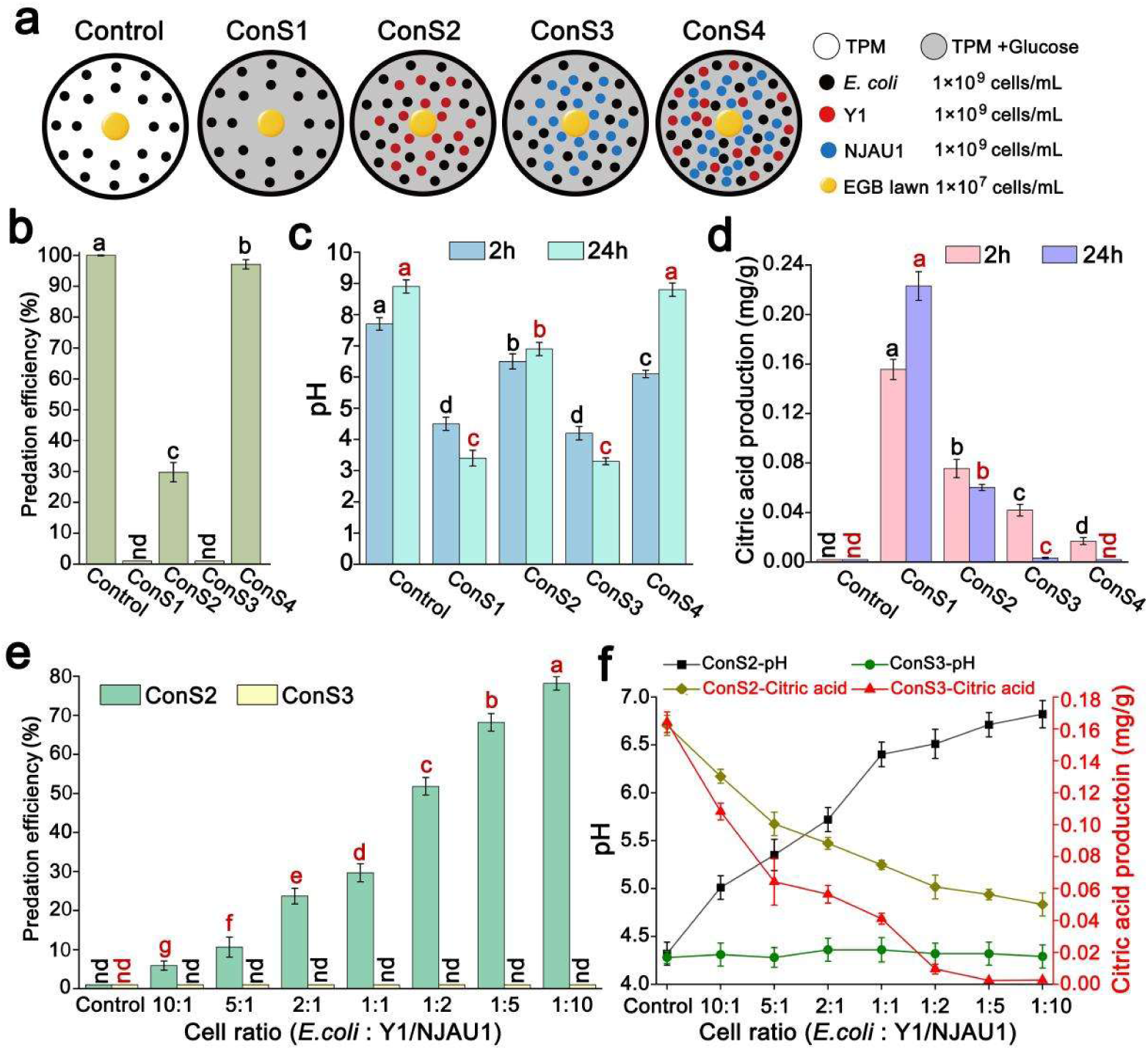
Soil microbes restore the predation of *E. coil* by EGB under glucose-rich conditions. **a)** Experimental scheme of four synthetic communities (42) based on the different combinations of strains on agar plates. *E. coli* was mixed with Y1 and/or NJAU1 in equal proportions, with a final density of 5×10⁹ cells/mL for each strain. EGB (1×10⁷ cells/mL, 3 µL) were inoculated to the center of ConS spots (35 µL) on glucose-contained TPM agar plates and documented after 2 days. Cells were harvested and biomass of *E. coli* during co-culture was quantified by qPCR. The predation efficiency of EGB against *E. coli* (**b**) and temporal changes in environmental pH (**c**) and citric acid levels (**d**) from the co-culture assays were analyzed. By altering the ratios of *E. coli* to Y1 in ConS2 and *E. coli* to NJAU1 in ConS3 from 10:1 to 1:10, the predation efficiency of EGB against *E. coli* measured (**e**). In panel (**f)** the pH changes were measure at 2 h while citric acid accumulation was measured at 12 h. Error bars denote ± standard error of the mean (SD), and different lowercase letters indicate statistically significant differences (*p* < 0.01) between groups based on Student’s t-test. “nd” indicates not detected.

We observed that Y1 and NJAU1 possess different abilities to utilize glucose and citrate. To further address the function allocation, we changed the synthetic mixture by modulating the abundance ratios of *E. coli* to Y1 (in ConS2) or NJAU1 (in ConS3) across a gradient ranging from 10:1 to 1:10. Increasing the Y1 population led to a proportional enhancement of EGB predation efficiency and corresponding rise in pH within the first hour (Figs. 5e-f), along with a decrease in citrate accumulation (Fig. 5f). However, even though the ratio of *E. coli* to Y1 reached 1:10, trace citrate still accumulated to 0.05 mg/g after 12 (Fig. 5f), indicating the primary role of Y1 in reducing environmental acidification by consuming glucose, yet at 2 h the pH value remained largely unchanged. However, when the ratio of *E. coli* to NJAU1 reached 1:2, NJAU1 in ConS3 reduced citrate levels to 0.008 mg/g, which was below the minimum effective concentration (0.1 mM, about 0.02 mg/g) (Fig. 5f), indicating the role of NJAU1 in restoring EGB predation by efficiently metabolizing accumulated citrate. These results suggest that in a soil environment, myxobacteria may overcomes glucose-induced predation inhibition through a metabolic utilization strategy from other soli microbes: *M. guilliermondii* mitigates early acidification through competitive glucose consumption, while *Xanthomonas* effectively metabolizes citrate, thus both factors support myxobacterial predation.

### Population density and sugar metabolism determine the competition outcome of EGB and *E. coli*

We next explored possible factors that affect the predatory interaction of EGB and *E. coli* with glucose from a population perspective. By varying the cell density of the predator EGB on TPM plate containing glucose, we found that EGB failed to prey on *E. coli* after 2 day across a broad density range (10⁷-10¹⁰ cells/mL) (Fig. S16a), indicating that EGB is unable to mitigate glucose-induced resistance by increasing their population density under nutrient-limited conditions. In contrast, EGB predation with glucose input was *E. coli* density dependent. Compared to predation inhibition at high *E. coli* densities (≥1×10^9^ cells/mL; *E. coli*: EGB ratio ≥ 1000:1), EGB preyed on *E. coli* in the presence of glucose at lower prey densities (≤1 × 10⁹ cells/mL; *E. coli*: EGB ratio ≤ 1000:1) (Fig. 6a, Fig. S16b). The deletion of *ptsI*-*ptsH* relieved density-dependent predation resistance of *E. coli* toward EGB (Fig. S16b). Given that the population density of *E. coli* was negatively correlated with EGB predation efficiency or pH values at 2 h (Fig. 6a) and positively correlated with citrate accumulation (Fig. 6a), we concluded that predation resistance through PTS-mediated glucose metabolism was prey density-dependent.

**Fig. 6.**
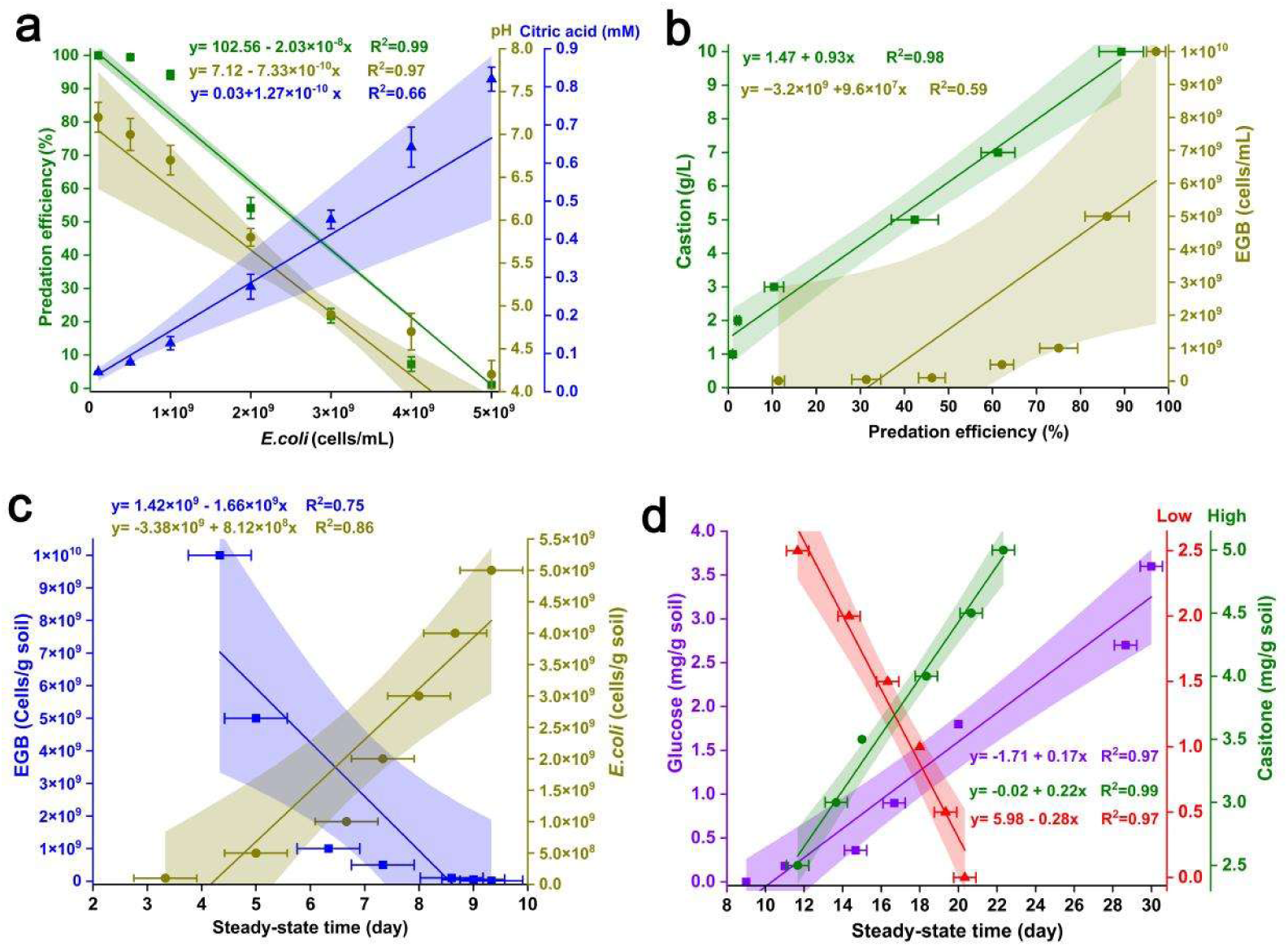
Population density and carbon-nitrogen input determine the predatory outcomes between EGB and *E. coli*. **a)** Correlations between the initial *E. coli* cell density and predation efficiency of EGB (green), the pH values (at 2 h, dark yellow), and citrate accumulation (at 12 h, blue). Predation efficiency of EGB on TPM agar plates containing 10 mM glucose by altering varying initial EGB and *E. coli* densities. EGB (1×10⁷ cells/mL, 3 μL) were inoculated to the center of *E. coli* spots (1×10⁸ to 5×10⁹ cells/mL, 35 μL), or EGB (1×10⁷ to 1×10¹⁰ cells/mL, 3 μL) were inoculated to the center of *E. coli* spots (5×10^9^ cells/mL, 35 μL) on agar plates containing 10 mM glucose and documented after 2 days. The biomass of residual *E. coli* was quantified using the plate colony counting method (CFU), and the predation efficiency of EGB was calculated accordingly. **b)** Correlations between casitone concentration and EGB predation efficiency (green), initial EGB cell density and predation efficiency (dark yellow) on TPM plates containing 3 g/L casitone. EGB (1×10⁷ cells/mL, 3 μL) were inoculated to the center of *E. coli* spots (5×10⁹ cells/mL, 35 μL) on TPM plates with 10 mM glucose and different concentrations of casitone (1-10 g/L), and the predation efficiency of EGB was determined after incubation at 30°C for 5 days. EGB and *E. coli* were co-cultured on TPM agar plates containing 10 mM glucose and 3 g/L casitone by altering varying initial EGB cell density. Following incubation at 30°C for 4 days, the predation efficiency of EGB against *E. coli* was measured. **c, d)** Correlations between steady-state time and initial cell densities of EGB (blue) or *E. coli* (dark yellow) (c), steady-state time and glucose (purple) or casitone (green for 2.5-5.0 mg/g; pink for 0-2.5 mg/g) concentrations (d) in mixed soil consisting of 10% non-sterile soil and 90% sterile soil. Steady-state time: time required to reach predation equilibrium—defined as the point at which *E. coli* biomass no longer exhibited significant change. Error bars denote ± standard error of the mean (SD), n=12. Correlation analysis was performed using SPSS 20.0 (IBM Inc., Chicago, IL, USA), and Pearson correlation coefficients were calculated to assess the relationships between variables.

Since previous research observed that the growth rate of *M. xanthus* grown in liquid culture increases depending on cell density (43), we therefore incubated co-cultures of EGB and *E. coli* populations (ratio of EGB and *E. coli*, 1:5,000) on glucose-contained TPM plates with different levels of casitone. The results showed that EGB fails to prey on *E. coli* on TPM agar plates with 1 g/L casitone and 10 mM glucose (Fig. 6b). However, when casitone supplement exceeded 2 g/L, EGB restored its predation by alleviating glucose-mediated inhibition in a casitone concentration-dependent manner (Fig. 6b). Moreover, when the initial population density of EGB reached 1×10^10^ cells/mL (1:5 ratio of EGB to *E. coli*) on TPM plates containing 3 g/L casitone and 10 mM glucose, the predation efficiency recovered to over 97% (Fig. 6b). Under this condition, predation efficiency exhibited an obvious positive correlation with both EGB population density and casitone concentration (Fig. 6b), indicating that population density, along with the carbon and nitrogen sources, jointly determine the competitive outcome between predator EGB and prey bacteria.

To further determine the variation of the predator and prey population under the influence of relevant factors, we co-cultured EGB and *E. coli* in gnotobiotic soil system (10% non-sterile + 90% sterile soil) by adjusting the initial cell density, concentration of glucose and casitone. Following a 30-day co-culture period in soil, the final population of both predator and prey in all treatments had reached a stable state, and the tested variations failed to alter the final population densities, but influenced the time required to reach population equilibrium (Fig. S17). Without glucose input, the time to reach equilibrium showed a positive correlation with input *E. coli* density, but a negative correlation with initial EGB density (Fig. 6c). In addition, glucose concentration positively correlated with the time to equilibrium, extending from 11 days at 0.18 mg/g to 30 days at 3.6 mg/g (Fig. 6d), reinforcing the conclusion that glucose availability bolsters defense of *E. coli* against predation. In contrast, we observed a diauxie phenomenon with additional casitone. At low concentrations of casitone in the 0-2.5 mg/g range, a negative correlation was found between casitone concentration and population equilibrium. While, at high concentrations of casitone in the 2.5-5.0 mg/g range, a positive correlation was found (Fig. 6d), indicating an additional factor affecting the predatory interactions between EGB and *E. coli*.

### EGB releases ammonia from amino acid metabolism to combat prey acidification

To investigate whether nitrogen input enhances EGB predation by mitigating glucose-mediated inhibition, we designed a co-culture experiment to monitor nitrogen metabolism. We showed that EGB fails to prey on *E. coli* in glucose-supplemented Aci-SUP1 medium, where prey-mediated glycolysis acidifies the culture environment (Fig. S3, Fig. S18a). Nevertheless, when alkalized supernatant collected from predators in vegetative monoculture or fed with various prey was added to the co-culture system, EGB against *E. coli* was partially restored (Fig. S18a). Under these conditions the pH values in the co-culture supernatant gradually increased (up to 9.1-9.2), but not when EGB lacked prey (Fig. S18b). Prior work demonstrated that *M. xanthus* vegetative cells use peptides and amino acids in casitone as sources of carbon and nitrogen (43), and ammonia produced by amino acids metabolism are known to induce a pH change (44). We therefore suspected EGB predation responds to prey environmental acidification by the secretion of metabolized amino acids, i.e. ammonia, resulting in a pH increase. We tested this *in vitro* using an ammonia detection method (Nessler’s reagent), and found that EGB accumulated 18.6 mM and 23.8 mM ammonia levels when co-cultured with *E. coli* in TPM or CTT, respectively; while only trace ammonia levels were detected from *E. coli* monoculture in CTT (2.46 mM) or TPM (0.35 mM) medium (Fig. S18c), consistent with the observed pH changes (Fig. S18b).

Volatile ammonia from bacteria disperses through physical barrier from a distance (44). To assess whether EGB can do this, we designed a partitioned petri plate assay using phenol red as an indicator, which changes from pale orange to bright pink when the media becomes more alkaline. EGB, *E. coli*, or their co-cultures were grown physically separated by a polystyrene barrier, where volatile ammonia can pass over the barrier. As predicted, an obvious pH increase was seen caused by volatile compounds produced by EGB when co-cultured with *E. coli* on TPM plate or incubated on CTT plate after 3 days, where the receiver side turned bright pink, but not when EGB was incubated on a TPM plate or when *E. coli* was incubated on CTT or TPM medium (Fig. 7a). To determine the ammonia concentrations in the medium receiving by supplier side, the agar on the receiver side was extracted. Co-culture of EGB with *E. coli* on TPM and CTT plates accumulated 0.30- and 0.38-mg/g ammonia, and culture of EGB on CTT plate also produced 0.28 mg/g ammonia; while no ammonia was detected from control cultures (Fig. 7a). Predation assays on the left side further linked the increased pH value to restored bacterial predation, alkalization mediated by volatile ammonia from right side alleviates glucose-mediated predation inhibition on the left side (Fig. 7a), and the corresponding predation efficiency of EGB increased to 10.3% and 17.4% (left) from co-culture of EGB and *E. coli* on TPM and CTT (right), and to 33.5% (left) from pure culture of EGB on CTT (right) (Fig. 7a).

**Fig. 7.**
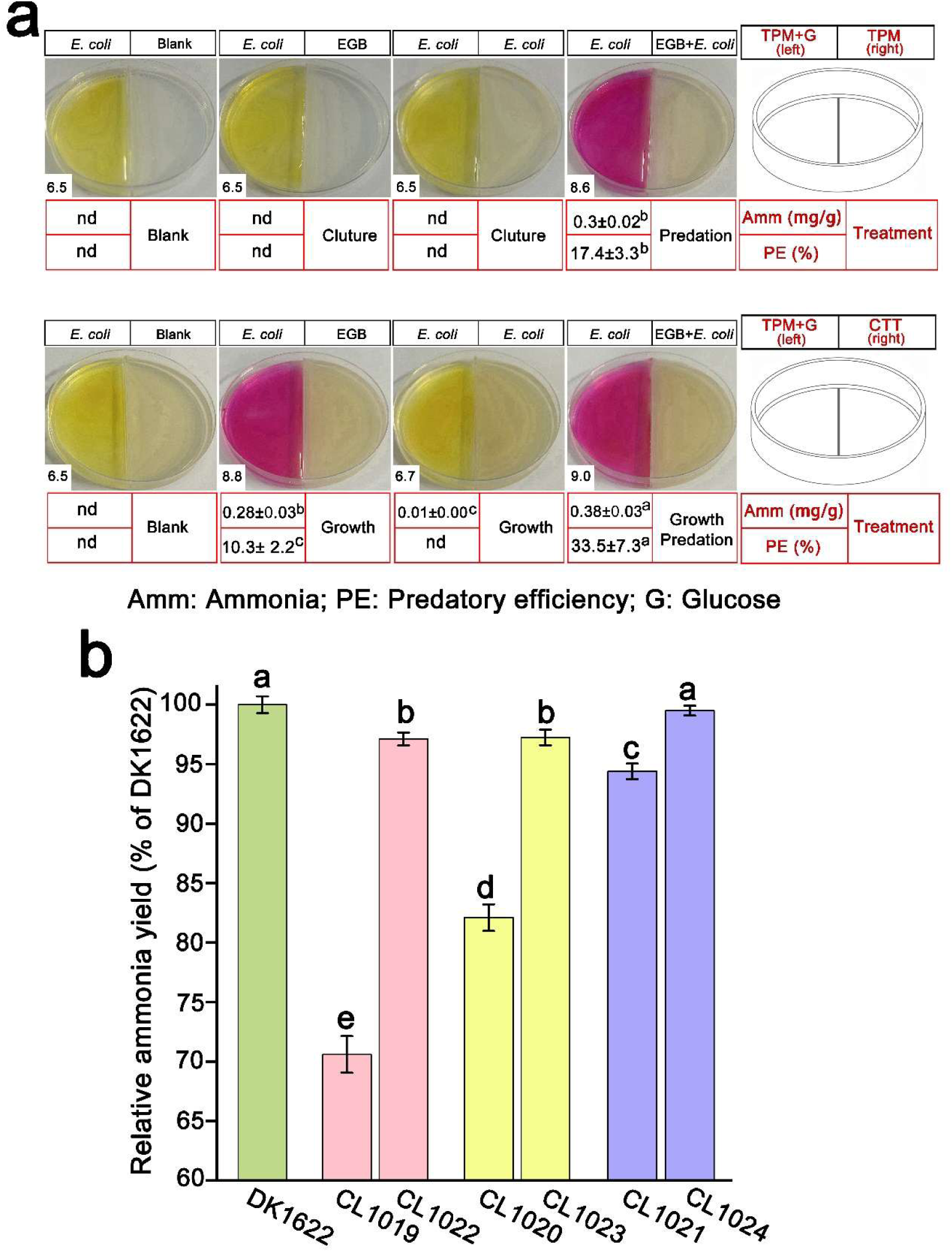
EGB produces volatile ammonia to counteract glucose-induced acidification. **a)** pH change illustrated by the color change of the indicator (phenol red, 0.002%) from split-plate assay (pH values shown in the lower-left corner). EGB and *E. coli* were cultured or co-cultured on right side of TPM and CTT agar plates, and left agar side was supplemented with phenol red when necessary. After incubation at 30 °C for 3 days, the ammonia level (Amm, mg/g) on the left side and predatory efficiency (PE, %) of EGB against *E. coli* on the right side were measured. **b)** Ammonia production by DK1622, amino acid cleavage mutants CL1019 (Δ*MXAN_1056*), CL1020 (Δ*MXAN_5198*), CL1021 (Δ*MXAN_2987*), and complementary strains CL1022 (Δ*MXAN_1056* :: *Kan-KYK13_RS04770*), CL1023 (Δ*MXAN_5198 :: Kan-KYK13_RS12135*), and CL1024 (Δ*MXAN_2987 :: Kan-KYK13_RS24610*). Using DK1622 as the reference (100%), the relative ammonia production of strains CL1019-CL1024 was determined in CTT medium after 1-day incubation. Error bars denote ± standard error of the mean (SD), and different lowercase letters (superscript in **a**) indicate statistically significant differences (*p* < 0.01, n=6) between groups based on Student’s t-test. “nd” indicates not detected.

We then wondered if ammonia was mainly generated from amino acid metabolism, a core energy source supporting vegetative growth of predatory myxobacteria. To evaluate which amino acid metabolism contributes the most to ammonia production, we designed the minimal TC3 medium and measured the released ammonia with the addition of various amino acids. Ammonia production by EGB significantly increased by 1.87-, 2.13- and 2.44-fold when co-cultured with Asn, IIe and Arg (Fig. S19a), respectively. A major pathway for the catabolism of arginine in *E. coli* is the AST pathway that converts arginine into ammonia and other metabolites (45). Additionally, prior research predicted that L-asparaginase and branched-chain keto acid dehydrogenase (BCKAD) are involved in catabolism of asparagine and isoleucine, respectively (27). Homologs corresponding to succinylarginine dihydrolase (KYK13_RS04770; MXAN_1056), L-asparaginase (KYK13_RS12135; MXAN_5198), and branched-chain amino acid aminotransferase (KYK13_RS24610; MXAN_2987) were identified in both EGB and the model organism *M. xanthus* DK1622. Upon cultivation with Arg, Asn and Ile, the expression levels of KYK13_RS04770, KYK13_RS12135 and KYK13_RS24610 were significantly upregulated by 8.0-, 4.8-, and 913.2-fold, respectively (Fig. S19b); this induction was similarly seen in DK1622 (Fig. S19b). Due to challenges in genetic manipulations of EGB, we instead deleted the homologous genes in the DK1622, generating mutants CL1019 (Δ*MXAN_1056*), CL1020 (Δ*MXAN_5198*), and CL1021

(Δ*MXAN_2987*). Ammonia production in CL1019, CL1020 and CL1021 decreased by 29.4%, 17.9% and 5.6%, respectively, compared to the wild type DK1622 (Fig. 7b). Complementation with the corresponding EGB genes (CL1022, CL1023 and CL1024) restored ammonia production to near-wild-type levels (Fig. 7b). These results demonstrate that EGB releases ammonia from amino acid metabolism to adapt the PTS-mediated extracellular acidification from prey.

### Ammonia derived from amino acid metabolism influences bacterial predator-prey interactions via a possible feeding strategy

We verified that EGB alkalinizes the surrounding environment by producing ammonia to relieve prey-induced acidification (Fig.7a). However, the long time- to-equilibrium at higher nitrogen concentrations (Fig. 6d) suggested the existence of an ecological mechanism in nitrogen metabolism that has not been recognized. Given that prey *E. coli* grew on media with ammonium (Fig. S20), we hypothesize that the metabolic by-products of ammonia from EGB predation or casitone metabolism could promote prey *E. coli* growth, thereby affecting the population equilibrium by a feeding behavior. To test this, predator EGB and prey *E. coli* (right) were grown physically separated from indicator *E. coli* by a polystyrene barrier on a TPM plate, where the ammonia can pass over the barrier, but it does not allow the passage of soluble nutrients (Fig. 8a). After 3 days, under the ammonia-emitting condition, co-culture of EGB and prey bacteria, caused population growth of all the indicator strains cultured adjacent to the supplier, while those inoculated next to the pure culture of EGB on TPM failed to grow (Fig. 8a). The enhanced growth of the indicator strain was accompanied by the consumption of the produced ammonia (Fig. 8a), indicating that in addition to neutralized acidification, EGB-derived ammonia also serves as a nitrogen source for prey. Furthermore, this regulatory phenomenon is prevalent among diverse soil bacteria (Fig. S21).

**Fig. 8.**
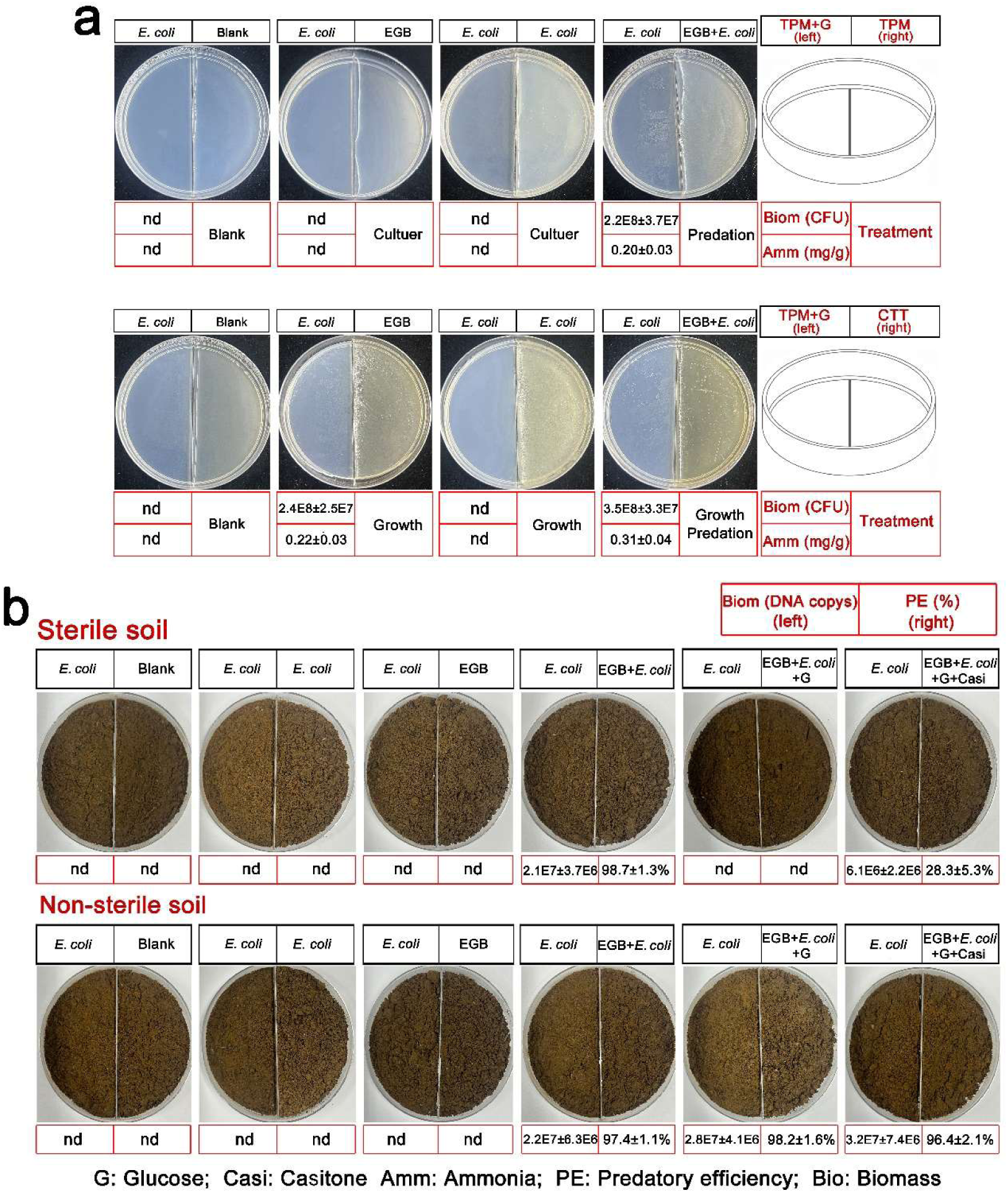
Growth promotion of bacteria by volatile ammonia produced by EGB. **a)** Cross-barrier volatile ammonia nutrient supply experiment (divided plate system). EGB and *E. coli* were co-cultured or monocultured on right side of TPM and CTT agar plates, pure *E. coli* (1×10^6^ cells/mL,100μL) spread on the left agar side was used as indicator. After incubation at 30 °C for 6 days, the consumption of ammonia and the increase in the biomass of *E. coli* on the left side were measured. **b)** Using *E. coli* as the prey and growth indicator, sterile or non-sterile soil was used to replace the medium in **a**). Two additional treatment groups were set up on the donor side (right): EGB + *E. coli* + 10 mM glucose, and EGB + *E. coli* + 10 mM glucose + 3 g/L tryptone. After 10 days of co-culture, soil samples were collected for DNA extraction, and the biomass of EGB and *E. coli* was determined by qPCR. Based on these data, the predation efficiency on the donor side and the biomass increase of *E. coli* on the left side were measured. “nd” Indicates not detected, n=6.

To examine whether ammonia-mediated trophic interactions between myxobacteria and prey bacteria also occur under more ecological conditions, we introduced EGB and *E. coli* into soil and assessed the growth-promoting effects of volatiles emitted from the soil (Fig. 8b). Without glucose input, EGB effectively preyed on *E. coli* in both sterile and non-sterile soils (Fig. 8b), and the biomass of indicator *E. coli* cultured adjacent to the supplier increased by 21-fold and 22-fold (Fig. 8b), while those inoculated next to the pure culture of EGB or *E. coli* failed to grow (Fig. 8b). However, glucose input inhibited EGB predation in sterile soil and thus failed to promote indicator growth (Figs. 8b), consistent with glucose-mediated predation inhibition (Fig. 2a-b). Nitrogen mentalism from casitone partially alleviated this inhibition, resulting in 6.1-fold increase in indicator biomass (Fig. 8b). Notably, in non-sterile soil, glucose did not impede EGB predation (Fig. 8b), and growth stimulation of adjacent indicator *E. coli* persisted (Fig. 8b), underscoring the role of soil microbes in mitigating inhibition. These findings revealed that the predator EGB exploits the ammonia generated from amino acid metabolism to support prey growth, maintaining a stable food source and thereby modulating the predator-prey population equilibrium.

## Discussion

Trophic interactions between microbial predators and prey are important mechanisms regulating soil food-web community structure and ecosystem function. Most studies on predation in soil microbial food web have focused on the predatory behaviors at higher trophic levels (46). In contrast, less attention has been given to bacterial predatory dynamics derived from metabolic interplay and resource utilization. As keystone taxa in the microbial food web, myxobacteria show promise for determining plant and soil health and recycling of soil nutrients (6, 24, 25, 47). Predatory myxobacteria are known to mainly metabolize amino acids from feeding on prey, but fail to utilize exogenous glycans released from the decomposition of prey cell walls (21, 27). The acquisition of sugars as energy sources enables many microbes to expand their nutritional capabilities, however this is not the case for myxobacteria species (28, 48), and the ecological significance behind these metabolic difference remains unclear. In this study, we show the predatory dynamics mediated by inherent carbon-nitrogen metabolism by predator myxobacteria and prey soil microbes. Commonly, prey bacteria utilize sugars for growth accompanied by extracellular acidification. Here, we propose that amino acids gained from prey, predatory myxobacteria maintain a stable prey population by a feeding strategy via allocation of available metabolic resources to soil microbes. Our findings offer valuable insights into possible trophic positioning and functional contribution of predatory myxobacteria in microbial food webs and microbial ecology in the context of predator-prey interactions from the perspective of metabolic evolution.

Soil microbes constitute multitrophic food webs. Nematoda and protists are key components of the microbial food loop responsible for carbon and nutrients recycling by feeding on bacteria (49). Myxobacteria prey on a broad range of bacteria in a contact-dependent manner (19, 25), representing a distinct trophic interaction among bacteria independent of protists and nematodes. Using bacteria predator-prey interactions between *Corallococcus* sp. EGB and soil microbes as a model, we discovered a resource difference between myxobacteria and their prey that are derived from inherent metabolic differences. Soil microbes frequently metabolize sugars from plant-matter decay or decomposition of macromolecules (50). In contrast, predatory myxobacteria utilize amino acids from prey lysis, while sugar metabolism by prey serves as a defense mechanism. Here we show that ammonia production from amino acid metabolism by EGB impacts soil microbes metabolite consumption that assists myxobacteria to adapt to sugar metabolism by prey. This complex metabolic interplay leads to predatory myxobacteria-prey bacteria oscillations until coexistence at equilibrium is reached, thereby constructing a relatively independent trophic interaction within a bacterial population. In another case, *Paenibacillus* exploits neighboring bacterial species to induces swarming away from a glucose-rich area, thereby override glucose-dependent swarming repression (51). Additionally, metabolism of the bacterial food glucose by other soil microbes could relieve glucose-mediated predation inhibition and assist myxobacteria to adapt to glucose-rich niches, thus increasing their fitness to various habitats.

Resource competition mediated by predation or defense is an important topic in ecological research. Microbes often modify their environmental pH to inhibit predators, like ciliate and *Bdellovibrio* (52, 53). Previous research has found that myxobacterial adaption to low pH and population growth are density-dependent (54, 55), but the underlying mechanism and its predatory significance remain unclear. Here we identified that EGB releases ammonia from deamination via inherent amino acid metabolism to adapt the PTS-mediated extracellular acidification from prey. In other interkingdom interactions, *Streptomycetes* kill bacteria competitors over long distances via ammonia release (44), and ammonia also regulate carbohydrate metabolism of mutualistic fungus *Leptographium procerum* of bark beetles to alleviate nutritional competition (56). This ammonia-mediated metabolic adaption by predatory myxobacteria represents another paradigm of inter-species’ interactions.

Common to other farming systems, bacterial farming by fungus has been observed from interaction between *Morchella crassipes* and *Pseudomonas putida* (16), where the fungus gains through the harvesting of carbon sources by bacterial farming via metabolic cross-talk. In addition to functioning in acid defense, we found that the by-product ammonia from amino acid metabolism of myxobacteria could support soil microbial growth, including prey bacteria. We propose that this feeding behavior provides a stable food source for predatory myxobacteria within the bacterial community. Nutrients release from cell lysis mediated by type VI secretion system (T6SS) and viruses participates in marine biogeochemical cycling, as proposed for the“bacterial shunt” and “viral shunt”, respectively (17, 57), and bacteriophage-driven bacterial killing may also contribute nutrients cycling that supports bacterial growth (58). Our analyses suggest that there may be an analogous “myxobacterial shunt” that participates in soil biogeochemical cycling.

In summary, our findings highlight the ecology significance of carbon-nitrogen metabolism in the interactions of predatory myxobacteria and soil bacteria, which provides experimental evidence for the role of myxobacteria as a keystone taxa influencing microbial population dynamics, and trophic positioning and functional contribution of predatory bacteria. Moreover, our results offer new insights into the possible metabolic evolution of predator and prey adaption, and whether such behaviors exist among myxobacteria in natural habitat is an intriguing but unexplored idea.

## Acknowledgements

We thank Daniel Wall (University of Wyoming) for helpful comments. This work was supported by the National Natural Science Foundation of China grants (32371730 to Z.C., 32270060 and 32570039 to Z.L., 32600157 to L.Z.), the National Natural Science Foundation of Jiangsu grants (BK20251549 to L.Z.), the Fundamental Research Funds for the Central Universities (QTPY2026009 to Z.L.), the Postdoctoral Fellowship Program of CPSF under Grant Number GZC20251872 (L.Z.), and Jiangsu Funding Program for Excellent Postdoctoral Talent under Grant Number 2025ZB725 (L.Z.).

## Reference

1. Dee LE, Ferraro PJ, Severen CN, Kimmel KA, Borer ET, Byrnes JE, Clark AT, Hautier Y, Hector A, Raynaud X. 2023. Clarifying the effect of biodiversity on productivity in natural ecosystems with longitudinal data and methods for causal inference. Nature Communications 14:2607.

2. Wall DH, Nielsen UN, Six J. 2015. Soil biodiversity and human health. Nature 528:69–76.

3. Sinclair A. 2003. Mammal population regulation, keystone processes and ecosystem dynamics. Philosophical Transactions of the Royal Society of London. Series B: Biological Sciences 358:1729–1740.

4. Thakur MP, Geisen S. 2019. Trophic regulations of the soil microbiome. Trends in microbiology 27:771–780.

5. Potapov AM. 2022. Multifunctionality of belowground food webs: resource, size and spatial energy channels. Biological Reviews 97:1691–1711.

6. Petters S, Groß V, Söllinger A, Pichler M, Reinhard A, Bengtsson MM, Urich T. 2021. The soil microbial food web revisited: Predatory myxobacteria as keystone taxa? The ISME Journal 15:2665–2675.

7. Wan B, Barnes AD, Jochum M, Chen X, Allen K, Yao J, He N, Hu F, Liu M. 2026. Aridity-induced energy reallocation from green to brown food webs predicts grassland carbon storage. Proceedings of the National Academy of Sciences 123:e2534463123.

8. Nguyen TBA, Chen QL, Yan ZZ, Li C, He JZ, Hu HW. 2023. Trophic interrelationships of bacteria are important for shaping soil protist communities. Environmental Microbiology Reports 15:298–307.

9. Falkowski PG, Fenchel T, Delong EF. 2008. The microbial engines that drive Earth’s biogeochemical cycles. Science 320:1034–1039.

10. Pérez J, Moraleda-Muñoz A, Marcos-Torres FJ, Muñoz-Dorado J. 2016. Bacterial predation: 75 years and counting! Environmental Microbiology 18:766–779.

11. Saleem M, Fetzer I, Harms H, Chatzinotas A. 2013. Diversity of protists and bacteria determines predation performance and stability. The ISME journal 7:1912–1921.

12. Summers JK, Kreft J-U. 2022. Predation strategies of the bacterium *Bdellovibrio bacteriovorus* result in overexploitation and bottlenecks. Applied and Environmental Microbiology 88:e01082–01021.

13. Nair RR, Vasse M, Wielgoss S, Sun L, Yu Y-TN, Velicer GJ. 2019. Bacterial predator-prey coevolution accelerates genome evolution and selects on virulence-associated prey defences. Nature Communications 10:4301.

14. Li G, Liu T, Xie W, Liu Z, Li H, Whalen JK, Jousset A, Wei Z. 2024. Metabolites limiting predator growth wane with prey biodiversity. Proceedings of the National Academy of Sciences 121:e2410210121.

15. Hiltunen T, Friman V-P, Kaitala V, Mappes J, Laakso J. 2012. Predation and resource fluctuations drive eco-evolutionary dynamics of a bacterial community. Acta Oecologica 38:77–83.

16. Pion M, Spangenberg JE, Simon A, Bindschedler S, Flury C, Chatelain A, Bshary R, Job D, Junier P. 2013. Bacterial farming by the fungus Morchella crassipes. Proceedings of the Royal Society B: Biological Sciences 280.

17. Stubbusch AK, Peaudecerf FJ, Lee KS, Paoli L, Schwartzman J, Stocker R, Basler M, Schubert OT, Ackermann M, Magnabosco C. 2025. Antagonism as a foraging strategy in microbial communities. Science 388:1214–1217.

18. Wang J, Wang J, Wu S, Zhang Z, Li Y. 2021. Global geographic diversity and distribution of the myxobacteria. Microbiology Spectrum 9:10.1128/spectrum. 00012–00021.

19. Herrou J, My L, Monteil CL, Bergot M, Jain R, Martinez E, Mignot T. 2025. Tad pili with adaptable tips mediate contact-dependent killing during bacterial predation. Nature Communications 16:4425.

20. Li Z, Ye X, Liu M, Xia C, Zhang L, Luo X, Wang T, Chen Y, Zhao Y, Qiao Y. 2019. A novel outer membrane β-1, 6-glucanase is deployed in the predation of fungi by myxobacteria. The ISME Journal 13:2223–2235.

21. Zhang L, Dong C, Wang J, Liu M, Wang J, Hu J, Liu L, Liu X, Xia C, Zhong L. 2023. Predation of oomycetes by myxobacteria via a specialized CAZyme system arising from adaptive evolution. The ISME Journal 17:1089–1103.

22. Liu L, Xu F, Lei J, Wang P, Zhang L, Wang J, Zhao J, Mao D, Ye X, Huang Y. 2023. Genome analysis of a plasmid-bearing myxobacterim Myxococcus sp. strain MxC21 with salt-tolerant property. Frontiers in Microbiology 14:1250602.

23. Wang J, Guo X, Fan Q, Liu L, Lin K, Shi R, He X, Ye X, Huang Y, Ji Y. 2025. A predator–prey interaction between myxobacteria and fungi mediated by outer membrane vesicles. Journal of Advanced Research.

24. Ye X, Li Z, Luo X, Wang W, Li Y, Li R, Zhang B, Qiao Y, Zhou J, Fan J. 2020. A predatory myxobacterium controls cucumber Fusarium wilt by regulating the soil microbial community. Microbiome 8:49.

25. Zhang L, Ye X, Zhang Z, Li Y, Cui Z, Wall D, Li Z. 2026. Bacterial predator-prey interactions and their potentials in combating plant pathogens. Microbiological Research:128559.

26. Wrótniak-Drzewiecka W, Brzezińska AJ, Dahm H, Ingle AP, Rai M. 2016. Current trends in myxobacteria research. Annals of microbiology 66:17–33.

27. Curtis PD, Shimkets LJ. 2007. Metabolic pathways relevant to predation, signaling, and development. Myxobacteria: multicellularity and differentiation:241–258.

28. Irschik H, Reichenbach H. 1985. An unusual pattern of carbohydrate utilization in *Corallococcus* (*Myxococcus*) *coralloides* (Myxobacterales). Archives of microbiology 142:40–44.

29. Görke B, Stülke J. 2008. Carbon catabolite repression in bacteria: many ways to make the most out of nutrients. Nature Reviews Microbiology 6:613–624.

30. Watson BF, Dworkin M. 1968. Comparative intermediary metabolism of vegetative cells and microcysts of Myxococcus xanthus. Journal of Bacteriology 96:1465–1473.

31. Xia C, Zhao Y, Zhang L, Li X, Cheng Y, Wang D, Xu C, Qi M, Wang J, Guo X. 2023. Myxobacteria restrain Phytophthora invasion by scavenging thiamine in soybean rhizosphere via outer membrane vesicle-secreted thiaminase I. Nature Communications 14:5646.

32. Liu Y, Wang J, Zhang Z, Wang F, Gong Y, Sheng D-h, Li Y-z. 2021. Two PAAR proteins with different C-terminal extended domains have distinct ecological functions in *Myxococcus xanthus*. Applied and Environmental Microbiology 87:e00080–00021.

33. Zhang X, Li Y, Li Y, Li C. 2025. Development of a high-barrier, antimicrobial, and pH-sensitive nanocomposite film based on gellan gum/sodium carboxymethylcellulose/sodium alginate with nano-SiO2 for strawberry preservation and monitoring. International Journal of Biological Macromolecules 298:139983.

34. Miller GL. 1959. Use of dinitrosalicylic acid reagent for determination of reducing sugar. Analytical Chemistry 31:426–428.

35. Shi L, Günther S, Hübschmann T, Wick LY, Harms H, Müller S. 2007. Limits of propidium iodide as a cell viability indicator for environmental bacteria. Cytometry Part A: the Journal of the International Society for Analytical Cytology 71:592–598.

36. Bustamante VH, Martínez-Flores I, Vlamakis HC, Zusman DR. 2004. Analysis of the Frz signal transduction system of *Myxococcus xanthus* shows the importance of the conserved C-terminal region of the cytoplasmic chemoreceptor FrzCD in sensing signals. Molecular Microbiology 53:1501–1513.

37. Skotnicka D, Smaldone GT, Petters T, Trampari E, Liang J, Kaever V, Malone JG, Singer M, Sogaard-Andersen L. 2016. A Minimal Threshold of c-di-GMP Is Essential for Fruiting Body Formation and Sporulation in *Myxococcus xanthus*. Plos Genetics 12.

38. Brückner R, Titgemeyer F. 2002. Carbon catabolite repression in bacteria: choice of the carbon source and autoregulatory limitation of sugar utilization. FEMS Microbiology Letters 209:141–148.

39. Solé M, Rius N, Lorén JG. 2000. Rapid extracellular acidification induced by glucose metabolism in non-proliferating cells of *Serratia marcescens*. International Microbiology 3:39–43.

40. Nagaoka S, Murata S, Kimura K, Mori T, Hojo K. 2010. Antimicrobial activity of sodium citrate against *Streptococcus pneumoniae* and several oral bacteria. Letters in Applied Microbiology 51:546–551.

41. Vaara M. 1992. Agents that increase the permeability of the outer membrane. Microbiological Reviews 56:395–411.

42. Feng S, Tan CH, Constancias F, Kohli GS, Cohen Y, Rice SA. 2017. Predation by *Bdellovibrio bacteriovorus* significantly reduces viability and alters the microbial community composition of activated sludge flocs and granules. FEMS Microbiology Ecology 93.

43. Rosenberg E, Keller KH, Dworkin M. 1977. Cell density-dependent growth of *Myxococcus xanthus* on casein. Journal of Bacteriology 129:770–777.

44. Avalos M, Garbeva P, Raaijmakers JM, van Wezel GP. 2020. Production of ammonia as a low-cost and long-distance antibiotic strategy by Streptomyces species. The ISME journal 14:569–583.

45. Schneider BL, Kiupakis AK, Reitzer LJ. 1998. Arginine catabolism and the arginine succinyltransferase pathway in *Escherichia coli*. Journal of Bacteriology 180:4278–4286.

46. Robinson JM, Barnes AD, Fickling N, Costin S, Sun X, Breed MF. 2024. Food webs in food webs: The micro–macro interplay of multilayered networks. Trends in Ecology & Evolution 39:913–922.

47. Li Z, Zhang L, Ye X, Huang Y, Ji Y, Li Y, Wall D, Cui Z. 2025. Myxobacteria: versatile cell factories of novel commercial enzymes for bio-manufacturing. Biotechnology Advances 82:108594.

48. Bretscher AP, Kaiser D. 1978. Nutrition of *Myxococcus xanthus*, a fruiting myxobacterium. Journal of Bacteriology 133:763–768.

49. Bonkowski M. 2004. Protozoa and plant growth: the microbial loop in soil revisited. New Phytologist 162:617–631.

50. Gunina A, Kuzyakov Y. 2015. Sugars in soil and sweets for microorganisms: review of origin, content, composition and fate. Soil Biology and Biochemistry 90:87–100.

51. Dharanishanthi V, Orgad A, Rotem N, Hagai E, Kerstnus-Banchik J, Ben-Ari J, Harig T, Ravella SR, Schulz S, Helman Y. 2021. Bacterial-induced pH shifts link individual cell physiology to macroscale collective behavior. Proceedings of the National Academy of Sciences 118:e2014346118.

52. Teng Z-J, Wang P, Chen X-L, Guillonneau R, Li C-Y, Zou S-B, Gong J, Xu K-W, Han L, Wang C. 2021. Acrylate protects a marine bacterium from grazing by a ciliate predator. Nature Microbiology 6:1351–1356.

53. Dashiff A, Keeling TG, Kadouri DE. 2011. Inhibition of predation by *Bdellovibrio bacteriovorus* and *Micavibrio aeruginosavorus* via host cell metabolic activity in the presence of carbohydrates. Applied and Environmental Microbiology 77:2224–2231.

54. Fiegna F, Pande S, Peitz H, Velicer G. 2023. Widespread density dependence of bacterial growth under acid stress. iScience 26.

55. Marshall RC, Whitworth DE. 2019. Is “Wolf-Pack” predation by antimicrobial bacteria cooperative? Cell behaviour and predatory mechanisms indicate profound selfishness, even when working alongside kin. Bioessays 41:1800247.

56. Fanghua L, Wickham Jacob D, Qingjie C, Min L. 2020. An invasive beetle-fungus complex is maintained by fungal nutritional-compensation mediated by bacterial volatiles.

57. Suttle CA. 2007. Marine viruses—major players in the global ecosystem. Nature reviews microbiology 5:801–812.

58. Suttle CA. 2005. Viruses in the sea. Nature 437:356–361.

